# Combined free energy calculation and machine learning methods for understanding ligand unbinding kinetics

**DOI:** 10.1101/2021.09.08.459492

**Authors:** Magd Badaoui, Pedro J Buigues, Dénes Berta, Gaurav M. Mandana, Hankang Gu, Tamás Földes, Callum J Dickson, Viktor Hornak, Mitsunori Kato, Carla Molteni, Simon Parsons, Edina Rosta

**Affiliations:** Department of Chemistry, King’s College London, London SE1 1DB, United Kingdom; Department of Physics and Astronomy, University College London, London WC1E 6BT, United Kingdom; Computer-Aided Drug Discovery, Global Discovery Chemistry, Novartis Institutes for BioMedical Research, 181 Massachusetts Avenue, Cambridge, Massachusetts 02139, USA; Department of Physics, King’s College London, London WC2R 2LS, United Kingdom; School of Computer Science, University of Lincoln, Lincoln LN6 7TS, United Kingdom

**Keywords:** Ligand-protein unbinding, molecular kinetics, CDK2, free energy calculations, machine Learning, Collective Variable selection, CV identification, feature selection methods, Transition State Analysis, MLTSA

## Abstract

The determination of drug residence times, which define the time an inhibitor is in complex with its target, is a fundamental part of the drug discovery process. Synthesis and experimental measurements of kinetic rate constants are, however, expensive, and time-consuming. In this work, we aimed to obtain drug residence times computationally. Furthermore, we propose a novel algorithm to identify molecular design objectives based on ligand unbinding kinetics. We designed an enhanced sampling technique to accurately predict the free energy profiles of the ligand unbinding process, focusing on the free energy barrier for unbinding. Our method first identifies unbinding paths determining a corresponding set of internal coordinates (IC) that form contacts between the protein and the ligand, it then iteratively updates these interactions during a series of biased molecular-dynamics (MD) simulations to reveal the ICs that are important for the whole of the unbinding process. Subsequently, we performed finite temperature string simulations to obtain the free energy barrier for unbinding using the set of ICs as a complex reaction coordinate. Importantly, we also aimed to enable further design of drugs focusing on improved residence times. To this end, we developed a supervised machine learning (ML) approach with inputs from unbiased “downhill” trajectories initiated near the transition state (TS) ensemble of the string unbinding path. We demonstrate that our ML method can identify key ligand-protein interactions driving the system through the TS. Some of the most important drugs for cancer treatment are kinase inhibitors. One of these kinase targets is Cyclin Dependent Kinase 2 (CDK2), an appealing target for anticancer drug development. Here, we tested our method using two different CDK2 inhibitors for potential further development of these compounds. We compared the free energy barriers obtained from our calculations with those observed in available experimental data. We highlighted important interactions at the distal ends of the ligands that can be targeted for improved residence times. Our method provides a new tool to determine unbinding rates, and to identify key structural features of the inhibitors that can be used as starting points for novel design strategies in drug discovery.

## I. INTRODUCTION

A recent paradigm shift in drug design highlighted the importance of long residence time as a key objective in addition to strong binding affinity.^1^ The residence time defines the timescale of the ligand bound in the binding pocket.^2, 3^ It is related to the overall rate of the unbinding process, which could consist of several steps. This therefore requires information about the corresponding high energy transition states and free energy barriers, which is challenging to obtain. Even if a drug interacts strongly with its target (high binding affinity), a short residence time can significantly reduce the efficacy of the drug.^4^ While many successful drugs have been discovered on the basis of high binding affinity alone, recent studies have shown that for drug efficacy, the kinetics of drug-receptor binding may be, in some targets, more important than affinity.^2^ The complexity of the drug-target dissociation may also involve several steps and complex pathways. Accordingly, promising hit candidates with high affinity were discarded for the next step of the drug discovery process due to their low residence time.^5,6^

A major challenge in drug discovery is finding a fast and reliable method to predict kinetics of ligand-protein interactions.^7^ Importantly, for experimental determination of ligand kinetics, ligands first need to be synthesized, which can be expensive and time-consuming even for a moderate number of compounds. Different experimental methods have been used to obtain kinetics of ligand-receptor unbinding, such as radioligand binding assays, fluorescence methods, chromatography, isothermal titration calorimetry (ITC), surface plasmon resonance (SPR) spectroscopy, and nuclear magnetic resonance (NMR) spectroscopy.^6,8^ Radioligand binding assays and fluorescence binding assays require binding with radiolabelled ligands, where they exploit the physical-chemical characteristics of the ligand between their free and complexed forms with the target. Several successful assays have been used to predict ligand-protein unbinding, for example fluorescence resonance energy transfer (FRET)^9^ or fluorescence correlation spectroscopy (FCS).^10^ These methods can suffer from interference (especially fluorescence), lack of accuracy for short residence times, and high cost/hazard in the case of radioligands.^11^ SPR is the most widely used assay to measure rate constants associated with (kon and koff) of ligand-receptor unbinding. The receptors are immobilized to a sensor that can distinguish the protein between its ligand-free form and its bound form. This method is label-free; however, the attachment of the protein to the probe may influence the activity of the protein, due to conformational changes.^11^ To offer a screening approach that alleviates these difficulties, various computational techniques have been proposed as alternatives to estimating the kinetics of unbinding events.^12,13^

Molecular dynamics (MD) is a powerful computational tool to understand at an atomistic level the behaviour of biological processes such as protein-ligand interactions.^14^ Unbiased MD simulations were successfully used in the initial stage of the drug discovery process, using either multiple independent relatively short simulations,^15^ or using specialized computer architecture, such as ANTON, where microsecond long simulations are readily accessible.^13^ However, due to the limited timescales typically accessible via MD simulations, it is often challenging to obtain sufficient statistical sampling required to calculate kinetic and thermodynamic properties accurately. Drug-protein unbinding processes occur on long timescales, typically ranging from millisecond to hours, depending on the nature and the strength of the interaction between the ligand and target. Some drugs, for example, Aclidinium, Deoxyconformycin, or Tiotropium, have a half-life of hours,^16^ requiring prohibitively long time scale simulations and highly demanding computer resources, therefore enhanced sampling methods are required.^17^

To accelerate the simulations and sample rare events, different enhanced sampling techniques have been proposed to predict free energy barriers and uncover the kinetics of biological events.^18,19^ These methods include free-energy perturbation,^20,21^ metadynamics (MetaD),^22,23^ temperature-accelerated MD (TAMD),^24^ steered MD (SMD),^25^ milestoning,^26^ umbrella sampling (US),^27^ replica exchange,^28^ scaled MD,^29^ smoothed potential MD,^30^ transition path sampling,^31^ τ-Random Acceleration Molecular Dynamics Simulations (τ-RAMD)^32^ and more recently a combination of enhanced MD with machine learning.^33–35^ For most of these methods, a key factor is the identification of a collective variable (CV), representing a physical pathway, that allows the calculation of the free energy profile.^36^ Hence, correct identification of appropriate CVs becomes a problem, with very few practical ways to build them properly.^37–39^ These methods have already been used for ligand unbinding: for example, MetaD was used to predict the ligand-protein unbinding of p38 MAP kinase bound to type II inhibitors,^40^ where depending on the set of CVs chosen, different values for *k_off_* were obtained, and the closest *k_off_* to the experimental data is still one order of magnitude lower. More recently, it was found that using a combination of MetaD and quantum mechanics/molecular mechanics (QM/MM) simulations, a more accurate prediction of the kinetics can be achieved.^41^ The residence times of Sunitinib and Sorafenib in complex with the human endothelial growth factor receptor 2 have been calculated using Steered Molecular Dynamics (SMD).^42^ SMD was also used to calculate the unbinding free energy profile for TAK-632 and PLX4720 bound to B-RAF.^43^ In both works, the ligands could be distinguished qualitatively to assess shorter, or longer residence times, however, the predicted free energy barriers for the unbinding were significantly lower than the experimental data.

To produce accurate free-energy profiles using biased simulations with many important degrees of freedom, we need to define an ideal set of CVs that map the full path of the reaction coordinate.^44,45^ Usually, the vectors that describe this manifold are selected based on *a priori* chemical/physical intuition, typically based on the initial binding pose of the ligand. The same set of CVs are then kept constant and used for the full simulation. Considering only CVs from an initial structure implies possibly neglecting essential interactions that occur during the unbinding process, thus significantly affecting the free energy calculation. Additionally, structures resolved by X-ray crystallography or cryo-EM may capture the system in metastable states, which do not always reflect appropriate conformers for ligand binding.

In this work, we introduce a novel enhanced sampling method to obtain accurate free energy barriers for ligand-protein unbinding. Unlike existing methods, we also propose a method that subsequently can identify key molecular features determining the unbinding kinetics. We suggest an iterative way of assigning our CVs during the unbinding trajectory and then use these CVs as the driving force to pull the ligand out from the pocket and to perform the sampling for accurate free energy calculations. Similarly to *e.g.* τ-RAMD (which, however does not provide a free energy profile), there is no need to *a priori* select CVs; these naturally arise from the unbinding trajectories that build a reliable path of unbinding taking the flexibility and dynamics of the system into consideration.

The CVs extracted from our trajectories sufficiently describe a full pathway for the unbinding process. Subsequently, we optimize this path in the space of the identified CVs to obtain a minimum free energy profile using the finite temperature string method.^46^ While different unbinding trajectories may lead to slightly different variations due to multiple local minima along the paths, we typically expect that the main transition state ensembles would be captured by all of these paths similarly after the convergence to the minimum free energy pathway. This is the main underlying assumption behind the finite temperature string method, which was proven to work very well even for complex systems.^47,48^ Our results accordingly show little variations in the unbinding free energy barriers using different starting pathways for free energy calculations.

In addition to determining unbinding rates, we also aim to identify key molecular descriptors that provide guidance for further design of drugs based on improved residence times. We propose a systematic approach to identify key low-dimensional sets of internal coordinates using machine learning (ML) approaches. Machine learning methods have been widely successful in multidimensional data driven problems, which are also applied to biomolecular simulations to determine key CVs.^49–51^ Here, we develop a novel approach making use of our obtained string unbinding pathway and, within that, the knowledge of the transition state (TS) ensemble. We explored two different ML methods in this study: Neural Networks (NN),^52^ which provide efficient training on complex high-dimensional data, and Gradient Boosting Decision Trees (GBDT),^53^ which allow straightforward evaluation of feature importances (FI).^54^ We generate unbiased “downhill” trajectories initiated at our TS, and used these to train a ML model which predicts the fate of binding or unbinding.

To test this approach on a simple analytical model system, we generated trajectory data using a collection of 1D model potentials, including one selected double-well potential. Our results demonstrate that our novel ML analysis can identify the key features correlated to this selected double-well potential to define the end states and thus can be used for key feature selection successfully. To demonstrate the applicability and accuracy of this approach on challenging complex biomolecular systems, we obtained free energy barriers for two ligands bound to CDK2 with PDB IDs of 3sw4 (*18K*) and 4fkw (*62K*) (***Figure 1***).^55^ Cyclin Dependent Kinase 2 (CDK2) is a crucial regulator in eukaryotic cell growth: deregulation of CDK2 has been associated with unscheduled cell proliferation resulting in cancer progression and aggressiveness.^56,57^ Selective inhibition of this protein makes it an appealing target in treating multiple tumours of specific genotypes.^58^ Several molecules are currently under clinical evaluation as CDK2 inhibitors for cancer treatment, such as AT759,^59^ AG-024322,^60^ Dinaciclib,^61^ Roniciclib,^62^ Milciclib.^63^ Furthermore, CDK2 is an ideal benchmark system with its relatively small size and well-documented kinetic data for the binding of a range of different molecules.^55^

**Figure 1.**
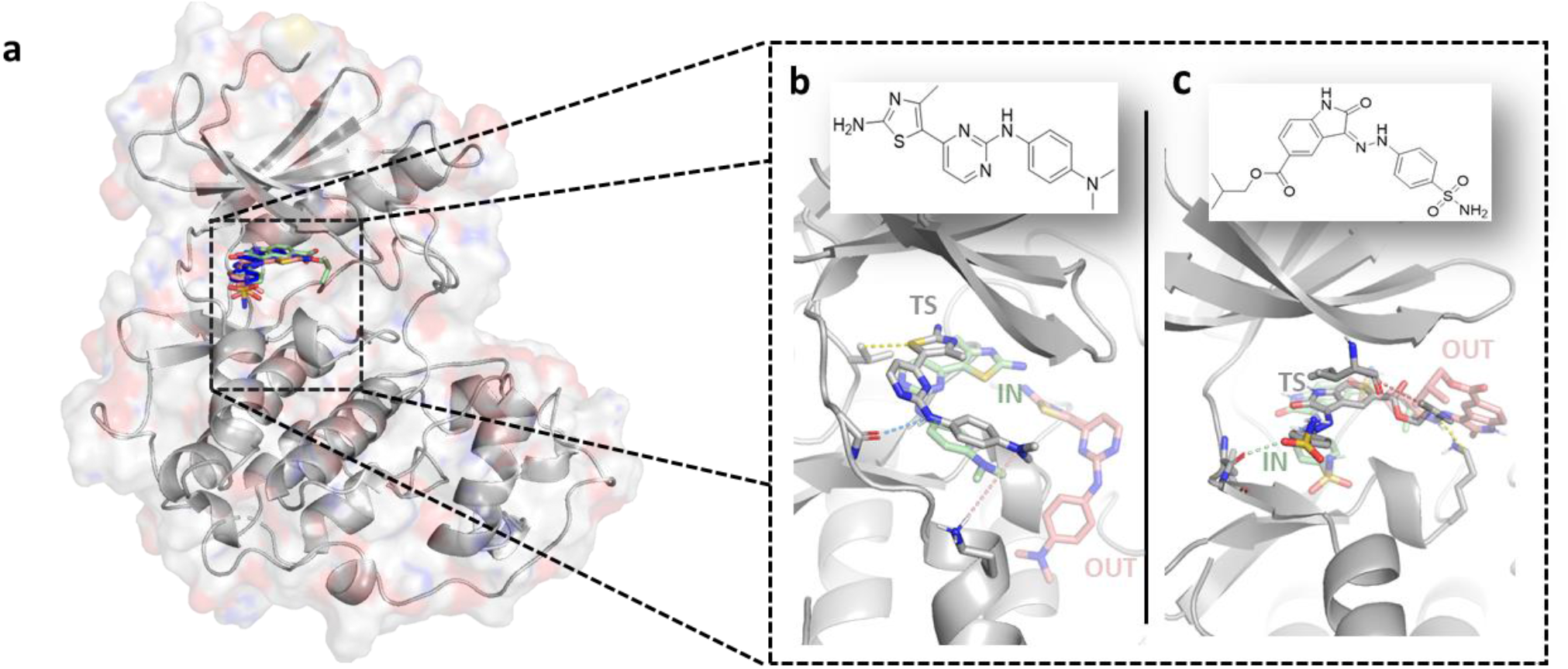
Illustration of the simulation system. **a** CDK2 bound to two different ligands: **b** thiazolyl-pyrimidine derivative (18K) and **c** carboxylate oxindole derivative (62K), originated from PDB structures 3sw4 and 4fkw, respectively. Structural details of the ATP pockets are shown for the two systems (bottom), with the ligands in the bound (green sticks), unbound (red sticks), and transition states (grey sticks). Dashed lines depict key interactions.

## II. METHODS

All MD simulations were carried out in NAMD 2.12, ^64^ using the AMBER ff14SB force field for the protein,^65^ and using the general Amber force field (GAFF) for the ligands.^66^ The MD simulation setup is detailed in SI Section 1.

### A. UNBINDING SIMULATIONS

Our unbinding method is illustrated algorithmically in ***Figure 2***. An explorational unbiased MD simulation of at least 20 ns was performed to identify the initial interactions between the protein and the ligand in the bound state. These initial simulations allow us to define the first set of CVs describing all distances between the heavy atoms of the ligand and the heavy atoms of the protein smaller than *d_in_* = 3.5 Å, our interaction cut-off. The identified interactions will generate a single one-dimensional CV as the sum of these *M* distances, *d*_*i*_, and will be used for iteratively biasing the simulations to observe an unbinding trajectory.

**Figure 2.**
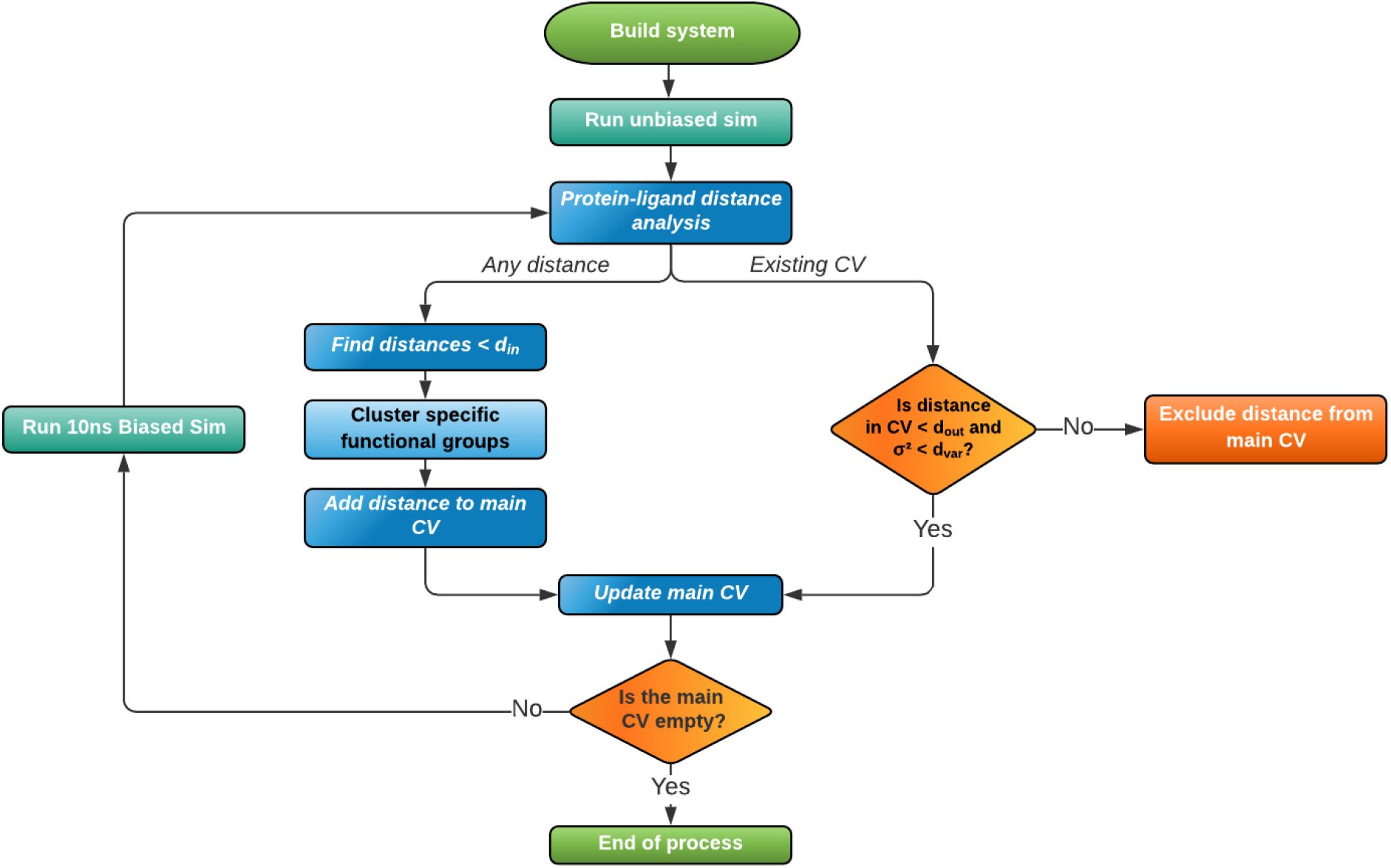
Flowchart illustrating the steps for the unbinding protocol.

At every iteration, we will define our bias as a harmonic restraint: 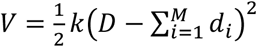, where *D* = *D*_0_ + (*Md*_*tar*_). Here, we aim to reach the target value *D* for our 1D CV starting from the initial value at the beginning of the *n*^th^ iteration *D*_0_. *d*_*tar*_ is the incremental factor, set to 1 Å, representing the average increase we aim to achieve per distance for the next iteration. The targeted *D* value will be reached progressively within the next 10 ns long MD simulation for every iteration. The force constant, *k*, was set to 20 kcal/mol/Å^2^.

At the end of each iteration, the biased trajectory was analyzed, and novel interactions were identified, within *d_in_* of the ligand, that are present for more than half of the total simulation time (*i.e.*, 5 ns). These novel interactions are then added to the list of interactions that define the main CV for the next iteration. Additionally, we also re-evaluate existing interactions. If a distance during the last 5 ns of the trajectory exceeds *d_out_* = 6 Å or its variance exceeds *d_var_* = 1 Å, then the distance is removed from the main CV in the next iteration. This exclusion factor will ensure that once a protein-ligand atom pair distance has exceeded *d_out_*, and therefore there is no significant interaction between these atoms, this interaction is no longer biased. Similarly, loosely interacting atom pairs have higher distance fluctuations, and thus the corresponding weak interaction does not need to be included in the bias.

To reduce the number of interactions between the ligand and the protein and to remove redundancies, we combine atoms that are part of an equivalent group where a rotational degree of freedom can interconvert the atoms from one to the other (for example, benzene ring or carboxylic groups, Fig. S1). Here, we considered the centre of mass of that functional group and not the individual atoms.

The iterative process will end when no more distances are present in the main CV from the last iteration *n*, thus there are no more stable interactions between the ligand and the protein, suggesting that the ligand is outside the binding pocket. Figs. S2.I-VI represent the distances included in the unbinding trajectories.

### B. FREE ENERGY CALCULATIONS

Once the ligand is outside of the binding pocket, to determine the minimum free energy path for the unbinding trajectory, we use the finite-temperature string method.^46^ The initial path and the full set of distances (CVs) are taken from the obtained unbinding trajectory.^46,67,68^ We extract these CV values for each interatomic distance along the initial unbinding path to construct the minimum free energy unbinding pathway iteratively, building a string of 100 windows in the coordinate space. For each window and each CV, we apply a position restrain equidistantly along the initial fitted string, using a force constant of 20 kcal/mol/Å^2^. We perform biased simulations using these restraints for a total time of 5 ns per window. From the obtained set of trajectories, a high-order (8) polynomial fitting is applied using the average values for each collective coordinate to build the subsequent set of refined CV positions. The procedure is carried out iteratively until the convergence of the free energy profiles and the pathway. This is verified by ensuring that the maximal change of each CV between subsequent iterations is below 7% (or 0.3 Å) from the previous iteration. By adding multiple overlapping biasing potentials along the dissociation pathways which are parametrized via the identified CVs, the string simulations can sufficiently sample the high dimensional path describing the full unbinding trajectory in detail. Finally, to obtain the corresponding Potential of Mean Force (PMF), we unbias the simulations using the binless implementation^46^ of the weighted histogram analysis method (WHAM).^62^

We note that our method does not aim to calculate binding free energies or kon rates. These would require simulations of a completely dissociated ligand and protein system, for which the string method is not an efficient algorithm. To this aim, routinely used efficient and accurate FEP^14,70^ calculations can be combined with our method to determine binding free energies and koff rates, respectively, from which the kon rates can be derived.

### C. MACHINE LEARNING TRANSITION STATE ANALYSIS (MLTSA)

We developed a Machine Learning Transition State Analysis (MLTSA) method to identify novel descriptors that determine the fate of a trajectory from the TS, which is applicable to unbinding simulations, but also suitable for other applications as a low-dimensional feature selection method for highly complex processes where a TS region is identified. In our case, the novel molecular interactions between the drug molecule and the protein for unbinding provide key signatures that determine the unbinding kinetics.

To test the validity of the MLTSA, we created an analytical model and compared the ability of two ML approaches to detect correlated features: a Multi-Layer Perceptron (MLP) architecture NN model and Gradient Boosting Decision Trees (GBDT), a common ML approach in feature selection.

The analytical model was based on using multidimensional trajectories generated via a set of one-dimensional (1D) free energy potentials (SI Section 5). Two types of potentials were used, both a set of single-well (SW) and double-well (DW) potentials. We used all but one of the DW potentials as “noise” and one of the DW potentials to define the outcome of the process, as the decisive coordinate to classify trajectories as “IN” or “OUT”. We generated trajectories using Langevin dynamics along 25 1D potentials. We used these trajectories to define 180 input features analogously to our observable CV-s by computing linear combinations of the original coordinates (SI Section 5). In our example, 11 of these 180 contained the selected DW potential with some non-zero coefficient (Tables S1.I and S1.II). We used these set of CVs to train the ML methods to predict the trajectory outcomes. Importantly, we aimed to identify the CVs that had the largest coefficients for our key selected DW potential.

We trained the MLP to analyze the model datasets of the downhill trajectories and predict their possible outcome from early on data, *i.e.*, at 30-60 steps of the downhill trajectory for the analytical model. The training was performed using the Scikit-learn library.^71^ We trained a simple model with an MLP Classifier architecture, using three main layers (input, hidden, and output) with as many input nodes as input features depending on the system of study (for the analytical model 180 were used, for CDK2 see Table S2.II), fully connected to a hidden layer with 100 hidden neurons and ending in an output layer with one output node each for IN or OUT classifications. The model was optimized using the Adam solver^72^ and using the ReLu^73^ function as an activation function for the hidden layer. The training was done with a learning rate of 0.001, iterating over data until convergence or upon reaching the maximum number of iterations (500 epochs). Convergence is determined by the tolerance and the number of epochs with no change in loss. When there are 10 consecutive epochs with less than 0.0001 improvement on the given loss, the training stops, and convergence is reached. The same parameters were used for both the analytical model and CDK2 data.

We also tested the GBDT model using the Scikit-learn library as a comparison to the MLP approach. This method provides feature importances (FI) that enable the ranking and identification of relevant features. We trained 500 decision stumps as weak learners for GBDT minimizing a logistic loss function, with a learning rate of 0.1. The criterion for the quality of the splits was the Friedman Mean Squared Error (MSE), with a minimum of 2 samples to split an internal node, and a minimum of 1 sample to be at a leaf node. The maximum depth of the individual regression estimators was 3, without a limit on the maximum number of features to consider as the best split, without maximum on leaf nodes and using a validation fraction of 0.1. The same parameters were used for both the analytical model system and the CDK2 simulations.

The flowchart of the MLTSA method is illustrated in Fig. S3. For the analytical model, we run 180 trajectories for the ML training and a separate validation set with 50 additional unseen trajectories. Following the flowchart, after labelling them as “IN” or “OUT” using the decisive coordinate, we created a dataset for the ML algorithms containing 180 features per frame (Fig. S4). We trained the ML models at different time frames (Fig. S5) to observe the evolution of the accuracy throughout the simulations. The accuracy and number of epochs used in the training are given in Table S3. This allows us to find a time range in the simulations where the classification problem is neither hard nor too trivial. Using this range, we trained the MLP model to analyze the importance of the features with our novel method. In a similar fashion to feature permutation^74,75^, or other model inspection techniques^76–78^, the MLTSA uses the Global Mean (GM) approach^77^, which swaps the value of each feature, one at a time with the mean value of the feature across all data used for training. This altered dataset is used for prediction again expecting to get the same accuracy as the training on non-correlated features and an accuracy drop on the correlated features, which depends on the level of correlation. For the comparison with GBDT and its FI, we trained the model on the same time and fetched the FI from the model to compare it with the accuracy drop analysis (Fig. S6).

For the application of the MLTSA on CDK2, first we identified the approximate TS location by selecting the last simulation frames from the highest energy five windows near the TS point of the obtained PMF. From each of these five starting coordinates, we then ran 50 independent unbiased MD simulations, each 5 ns long. We classified and labelled these short ‘downhill’ trajectories by considering a combination of two key distances (Table S2.I), to identify which simulations finish either in a ligand bound position (IN) or in a ligand unbound position (OUT). We then selected the starting structure (i.e., our TS) that provides the closest to a 1:1 ratio of IN and OUT events amongst these trajectories, and we ran 200 additional 5 ns-long unbiased MD simulations with this starting point. We considered all interatomic distances (heavy atoms only) between the ligand and the protein within 6 Å at the TS starting position and determined the values of these distances along downhill trajectories (Table S2.II). These constitute a dataset of distances for each simulation trajectory, and we aimed to select the most important features from these with our MLTSA method.

The number of epochs and convergence of the loss function for each model can be found in Tables S4.I – S4.II and Fig. S7. Thus, using the frames coming from the multiple short unbiased MD simulation trajectories starting from our TS, we provided a dataset of distances extracted along the trajectory, as well as the future outcome of the IN or OUT events as the desired answer/classification. We performed the ML training at several different time ranges of the trajectories (Fig. S8), to observe the predicted accuracy at different time ranges along the simulations. From all the available trajectories for each system we reserve a part for further validation to avoid the overfitting of our model. The rest is used for training, with all frames from the trajectories concatenated and randomly mixed, then split in different fractions as training (0.7) and test (0.3) sets.. The trained model is additionally verified to have a similar prediction accuracy on the unseen trajectories.

Using our trained model, we assess which features are the most important for the model to predict whether the simulation is classified as bound (IN) or unbound (OUT). To do so, we apply our own feature reduction approach (FR), in which every single distance (i.e., feature) is excluded one-by-one from the analysis, and we calculate the drop in accuracy compared to the full set of distances present. Differently from the standard approach,^66^ where the real value of each excluded feature is replaced with a zero, here we replace the value for each excluded feature with the global mean of that selected feature across the simulations, thus cancelling the variance of the aforementioned feature.

## III. RESULTS AND DISCUSSIONS

### MLTSA analytical test and validation

ML training on the model potential-derived trajectories was performed with both MLP and GBDT ML methods. We performed the MLP training at different time frames and trajectory lengths, from the 0^th^ time step to the 500^th^ step in intervals ranging from 10 to 150 frames at a time to assess the accuracy through time (Fig. S5). Using a suitable time range consisting of the 30th-60th simulation steps from each trajectory, the trained ML methods found the classification problem accurately solvable, but not too trivial. We replicated the complete process 100 times by generating 180 new independent simulations for each replica and performing the ML training. The MLP achieved an average test accuracy of over 94% and an average validation accuracy of over 93%, whereas the GBDT achieved over 99% on the test set and 91% on the validation set.

To identify the selected DW potential and its highest correlated features from the dataset, we calculated the accuracy drop (MLTSA as in Fig. S3) using the trained MLP and compared this approach to the FI using GBDT. Training accuracies for both ML models at 1DW and 5DW potentials can be found in Table S3. Results of both feature analysis methods are found in Fig. 3 for the 1DW dataset and in Fig. S6 for the 5 DW potentials dataset.

**Figure 3.**
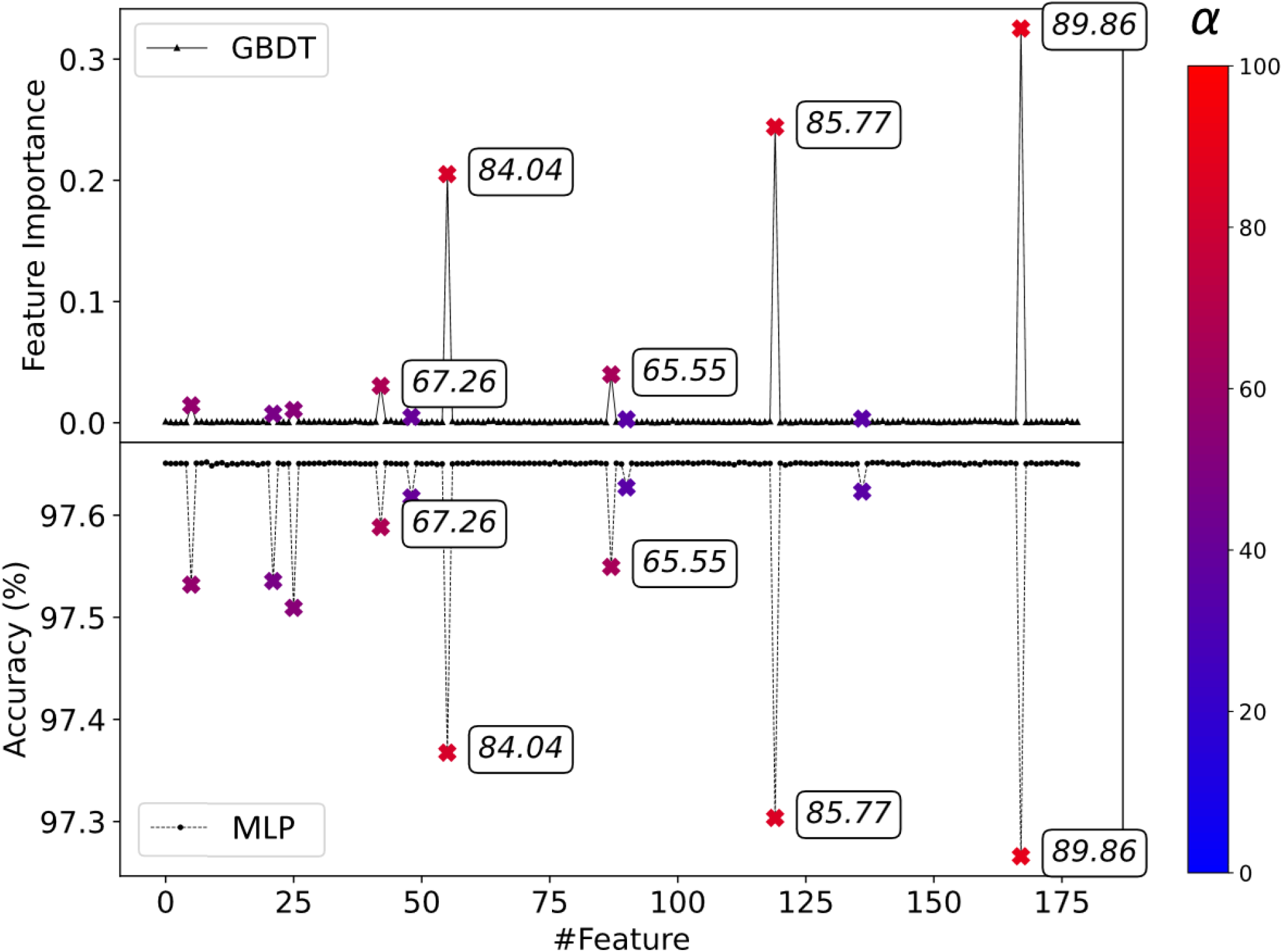
Comparison between GBDT (top) and MLTSA with NN (bottom) feature analysis methods for the 1DW dataset. Correlated features are marked from blue (0%) to red (100%) depending on the mixing coefficient, α (x symbols, color scale on the right, five highest mixing coefficients also displayed for the datapoints). Uncorrelated features (small black symbols) are at 0 FI for GBDT and show no loss of accuracy for MLTSA with MLPs. Correlated features all show a significant accuracy drop for the MLP, while only the top correlated features have high FI using GBDT.

The highest correlated features (colored depending on the correlation level, color bar in Fig. 3 right panel) were correctly identified by both MLP and GBDT models. For GBDT, only the top three features show a high FI value (labels added to datapoints in Fig. 3), whereas the rest of the correlated features ranging from α∼34% up to ∼60% do not show a significant FI value. In addition, despite three features (#48, #89 and #136) having 40.34%, 34.80% and 35.48% mixing coefficients, respectively, GBDT did not capture their correlation, showing values very close to 0. For the MLP, the top three distances are similarly captured as in the FI with the highest accuracy drops. Importantly, all correlated features have a non-zero accuracy drop, showing that they are correctly identified.

Using the dataset with increased complexity consisting of 5 DW potentials and 15 correlated features, we observed a similar performance of the two ML methods (Fig. S6). GBDT correctly captured and ranked the top three features (#8, #25 and #35). However, most other important features scored a FI value very close to 0. Out of 15 correlated features, GBDT did not identify 12 of them with high FI, whereas the MLP captured all of them. However, the MLP accuracy drop did not rank the top four features in the correct order, scoring the 3^rd^ most correlated feature with the biggest accuracy drop.

Considering both analytical models, we found that whereas GBDT has a higher specificity to rank the top correlated features in the correct order, MLP has a higher sensitivity and captures all correlated features but cannot necessarily identify the highest ranked ones quantitatively using the accuracy drop as the measure. Therefore, a combination of the two ML methods can further help identify the most important features. In more complex systems, this performance might not be directly generalizable, however, due to the simple linear correlation of the CVs of this model.

### CDK kinase unbinding free energy calculations

For each system, we performed three independent simulation replicas starting from the respective equilibrated system. For each replica, we performed the initial unbiased MD simulation, followed by our unbinding trajectory determination procedure and subsequently calculated the minimum free energy path and the corresponding free energy profile using the finite temperature string method (Fig. 4).

**Figure 4.**
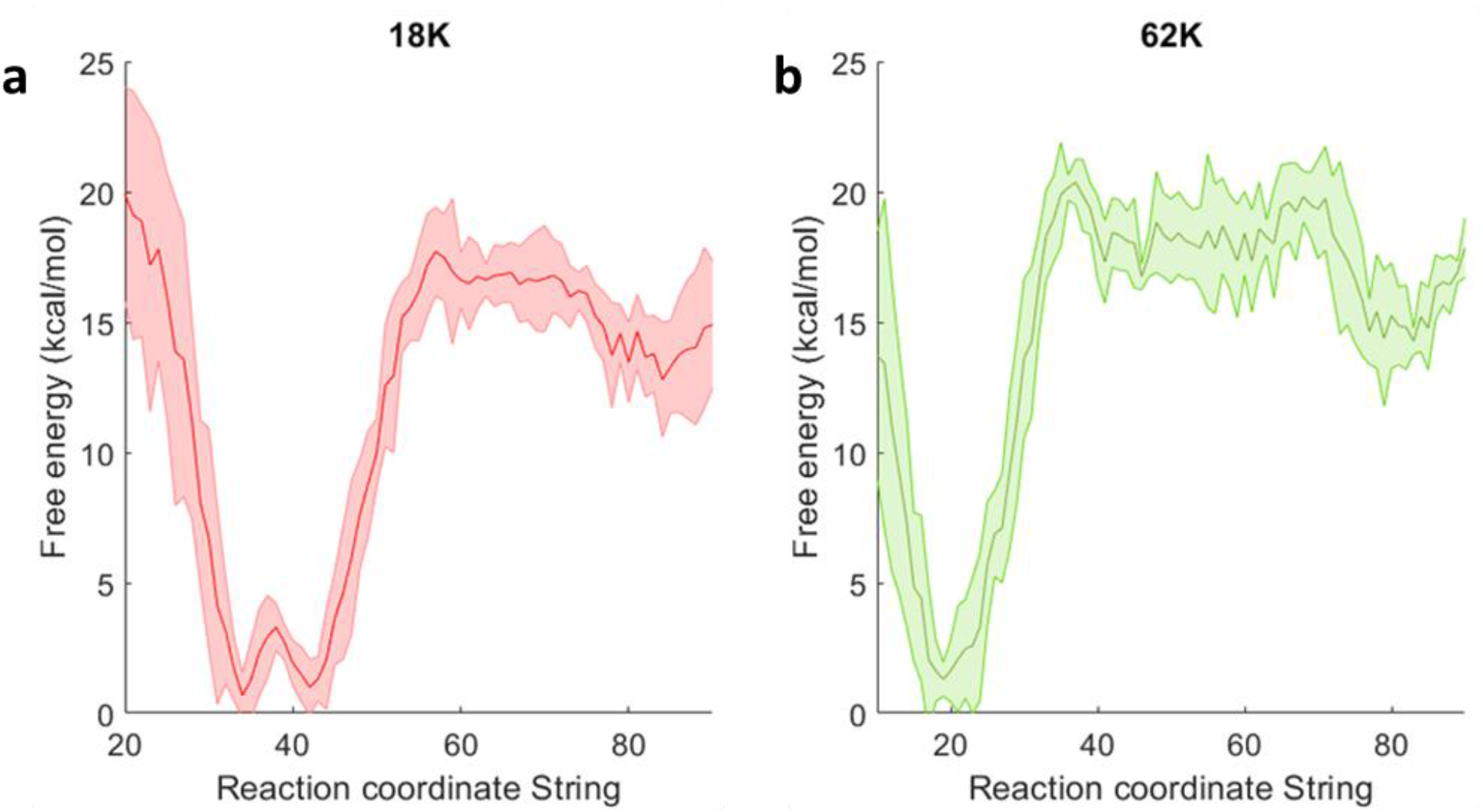
PMF of the unbinding path for 18K (**a**) and 62K (**b**). The free energy profile is obtained from a representative replica, the standard error, shown as a shaded area, obtained by dividing the full dataset into 4 subgroups.

Fig. 5 shows a representative result of the unbinding process for selected interactions. The first distance (blue line) is identified from the initial unbiased bound simulation as being shorter than 3.5 Å. Later during the biased unbinding process at 30 ns a new interaction is found (orange line) and at 90, 120 and 130 ns more distances are included in the main CV (green, red, purple, and brown). Additionally, interactions are progressively being removed as they are breaking (above 6 Å). Details of the selected CVs during the unbinding iterations are in the panels of Figs. S2.I-VI for every replica.

**Figure 5.**
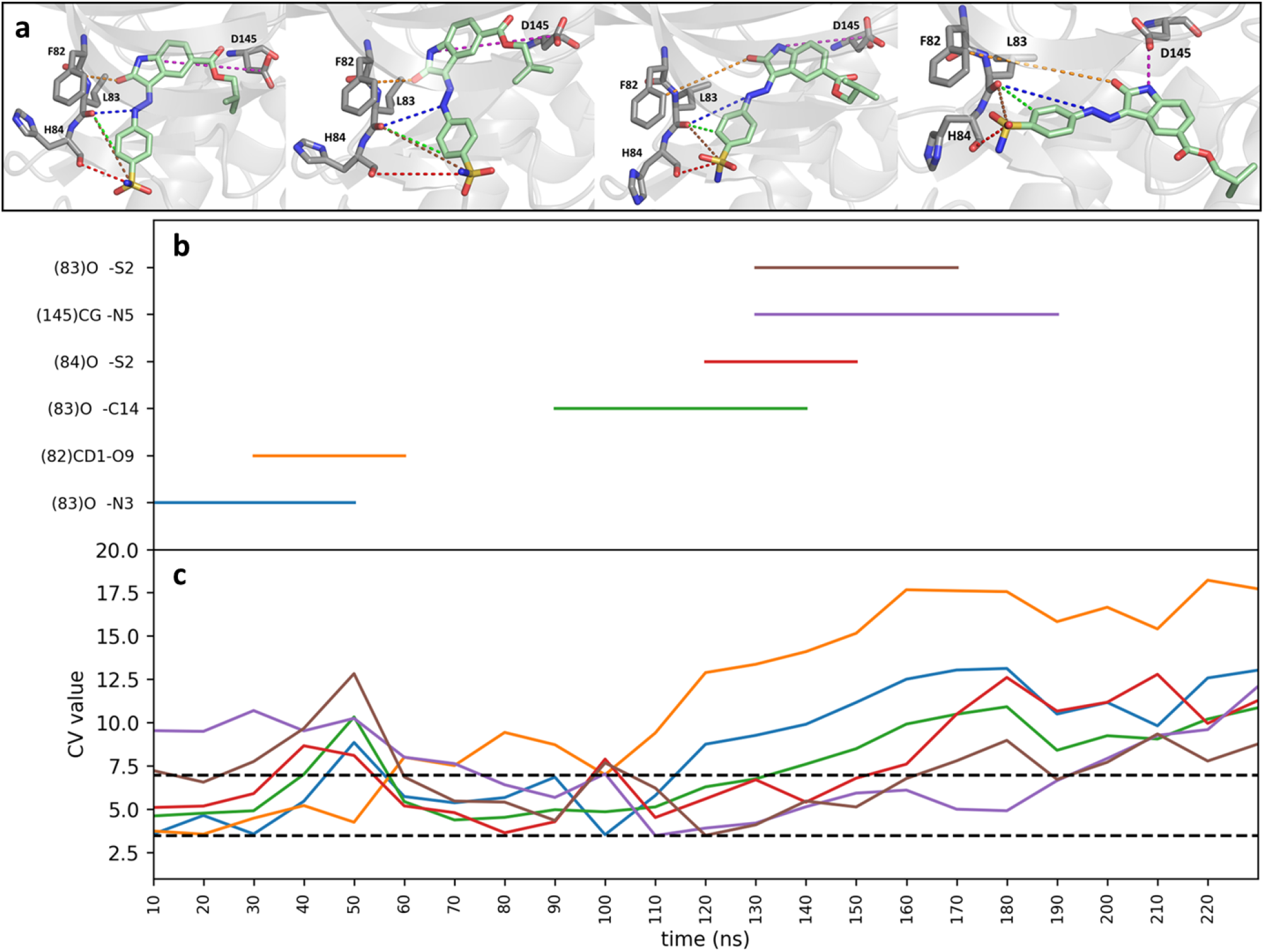
**a**) Unbinding trajectory of ligand 62K represented as selected snapshots along the trajectory at 0, 70, 141, and 219 ns from left to right, respectively. Representative distances used for the bias are shown as colored dashed lines (for the full set of distances please refer to Figs. S2.I-VI). Representative distances included in the CV along the unbinding trajectory are shown in (**b**), the corresponding distance values plotted in (**c**). The lower dashed line at 3.5 Å is the cut-off value below which an interaction is included in the main CV, the upper cut-off at 6 Å is the value above which the distance is excluded from the CV.

Overall, while the identified CVs in different replicas vary, a few common key CVs are present in all unbinding trajectories within all replicas (Fig. 6). Even if the actual unbinding pathways have differences amongst the replicas, as seen by looking at the distances found along the paths (Figs. S2.I-VI), they are all expected to pass through the same TS ensemble and show generally the same mechanism (see animated gifs for the final minimum free energy paths, SI Section 11 and SI Section 8 – 60K/4FKU system). This can also be confirmed from the consistent free energy profiles (Fig. S9).

**Figure 6.**
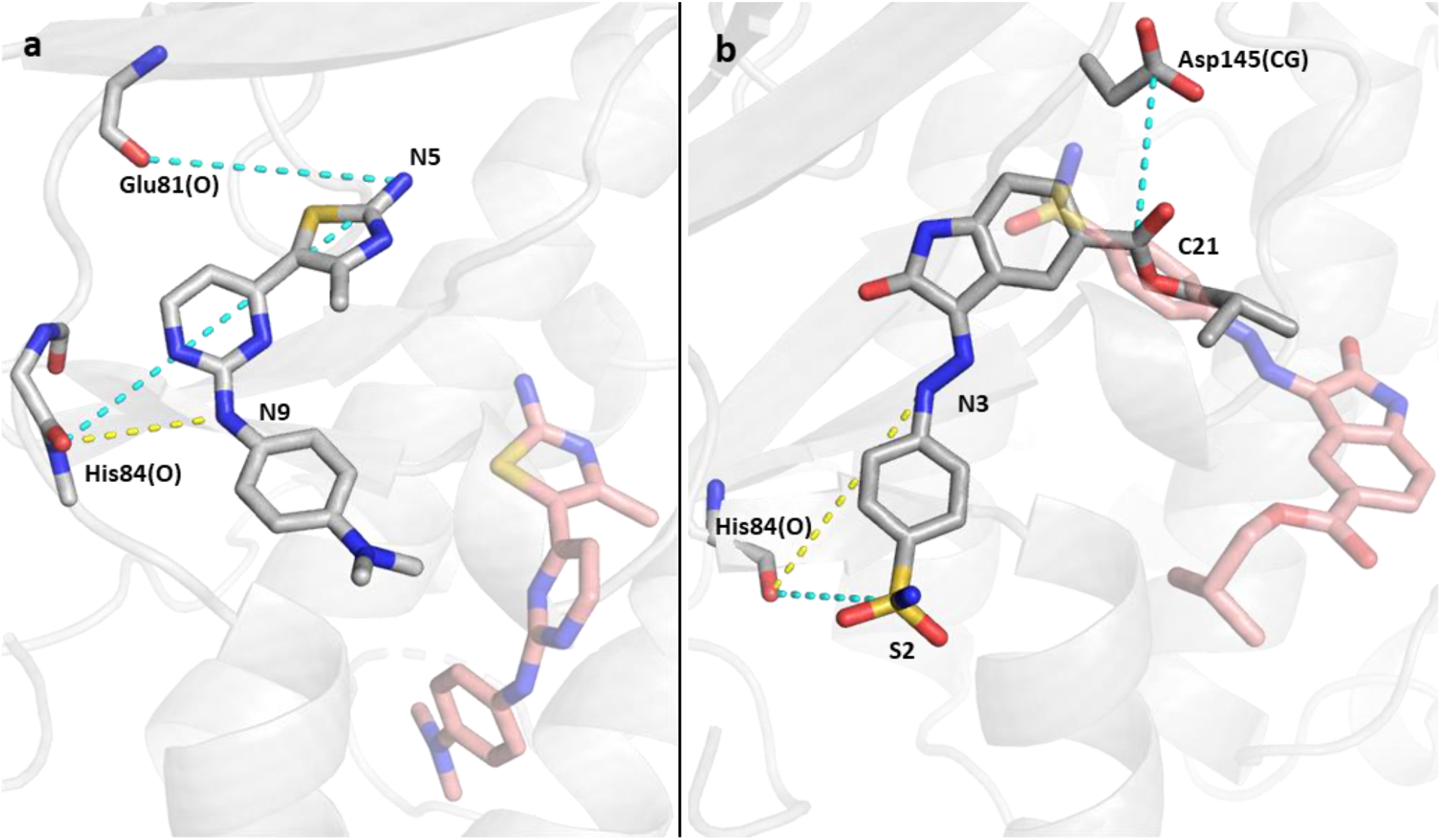
CVs obtained from the unbinding of 18K (**a**), and 62K (**b**); representative distances shown in dashed lines (yellow: interaction from the initial structure, cyan: interaction found during the unbinding trajectory), red sticks represent the coordinate of the ligand when it is outside the pocket. These distances appear in each of the three replicas for each system.

Additionally, we also performed the unbinding calculations for a third ligand, 60K, that is analogous to 62K (Fig. S10). Interestingly, we identified that all three replicate string pathways originating from three distinct unbinding simulations present a rotation of the hydrazineyl N=C bond, leading to a cis(Z)-trans(E) isomerisation of the ligand near the TS (Figs. S11 and S12). This is due to, on one hand, the initial strong forces in the string simulations that could be avoided in the future, and, on the other hand, to force field inaccuracies with a too low energy of the transform and too low barrier for the related dihedral angle rotation as determined by DFT calculations (Fig. S13). When compared with 62K, which does not exhibit this behavior in any of the three replicas, we can observe a lower energy for the 60K trans state, that enables it to avoid the TS bottleneck. Correspondingly, all three distinct replicas result in a consistently too low unbinding free energy barrier when compared with experiment (Fig. S9). Animated trajectories along the string simulations for all replicas are provided and can be accessed through SI Section 11 – Additional resources.

The energy barrier extracted from the PMF of our simulations agrees closely with the experimental koff rates and are very well reproducible within the same system (Table I and Fig. S9). The shape of the free energy profiles is also consistent amongst the replicas, however the exact shape depends on the CVs identified in that replica (Fig. S9 and Table S5). Generally, a higher number of CVs results in a broader TS region (e.g., Fig. S9, ligand 62K). In addition, results for the third ligand, 60K is also presented, demonstrating a consistent underestimation of the free energy barrier due to the discontinuity of the dihedral angle along the minimum free energy paths.^80^

**Table I.** Ligand binding kinetic and thermodynamic values of 3sw4 and 4fkw systems from Dunbar et al.^55^ and calculated results obtained from our simulations. ΔG^‡^_calc_ was calculated using the Eyring-Polanyi equation: k=k_B_T/h exp(-ΔG^‡^/k_B_T) at 298 K.^81^

| PDB | Ligand | $K_D$<br>(M) | $k_{\text{on}}$<br>[M<sup>-1</sup>s<sup>-1</sup>] | $k_{\text{off}}$<br>[s<sup>-1</sup>] | $\Delta G^{\ddagger}_{\text{exp}}$<br>(kcal/mol) | $\Delta G^{\ddagger}_{\text{calc}}$<br>(kcal/mol) |
| --- | --- | --- | --- | --- | --- | --- |
| **3sw4** | 18K | 9.61E-07 | 1.00E+05 | 0.0823 | 18.93(±0.17) | **16.29(±0.21)** |
| **4fkw** | 62K | 4.73E-08 | 6.49E+04 | 0.00261 | 20.97(±0.05) | **20.27(±1.06)** |

Importantly, comparing the same ligand within the three different replicas in all systems provide very similar free energy barriers, expressed with a low standard error. Our energy barriers consistently reproduce the high energy barriers also seen experimentally thanks to the introduction of numerous key CVs that are not only taken from the initial ligand-bound conformation but, instead, introduced along the unbinding paths (Fig. 5).

We observe only one main barrier corresponding to the breaking of the drugs with the His84 H-bonding contact (Fig. 4),^68^ suggesting that the different replicas do indeed share the same TS ensemble, despite the slightly varying pathways and identified CVs along the path. This H-bond was reported as a key interaction in many ligands in complex with CDK2/CDK5.^82,83^ This interaction was included in the initial unbiased simulation in the bound systems at the beginning of the unbinding procedure. However, during the unbinding trajectories, once this important H-bond between His84 and the ligand is broken, new interactions are formed, for varying time scales. For 18K, in all the three replicas, H-bonds are formed with the exocyclic amino group of the ligand (N5) and the backbone oxygen of Glu81 and subsequently with the backbone oxygen of His84. 62K presents a sulphonamide terminal group, which, during the trajectory, interacts with Val163 and His84 of CDK2.

To analyze which distances are the most important at the TS region, we implemented our MLTSA method. Starting with two datasets of 139 (62K) and 148 (18K) independent downhill trajectories for each system, and initial set of CVs of over 170 (Table S2.II), we obtained key distances for each system that are major determinants for the prediction of whether a molecule ends up in the bound or unbound states (Fig. 7). By training with trajectory data from up to 0.3 ns of each downhill simulation, the model can predict with high accuracy the IN or OUT outcome of the trajectories, more specifically: 80.11% for 18K and 93.83% for 62K. To confirm the effectiveness of the ML training, we compared the ML prediction accuracy with using optimal thresholds of our main string CVs (Fig. 7) to determine the outcome at 5 ns of downhill simulations (Figs. S14.I-II). Importantly, the ML model predicts the outcome more accurately at early times (before ∼0.3 ns), than using the best possible prediction via the string reaction coordinate: with above around 80% to 94% accuracy versus ∼55-to 61%, respectively for the ML and the main CV (Figs. S14.I-II).

**Figure 7.**
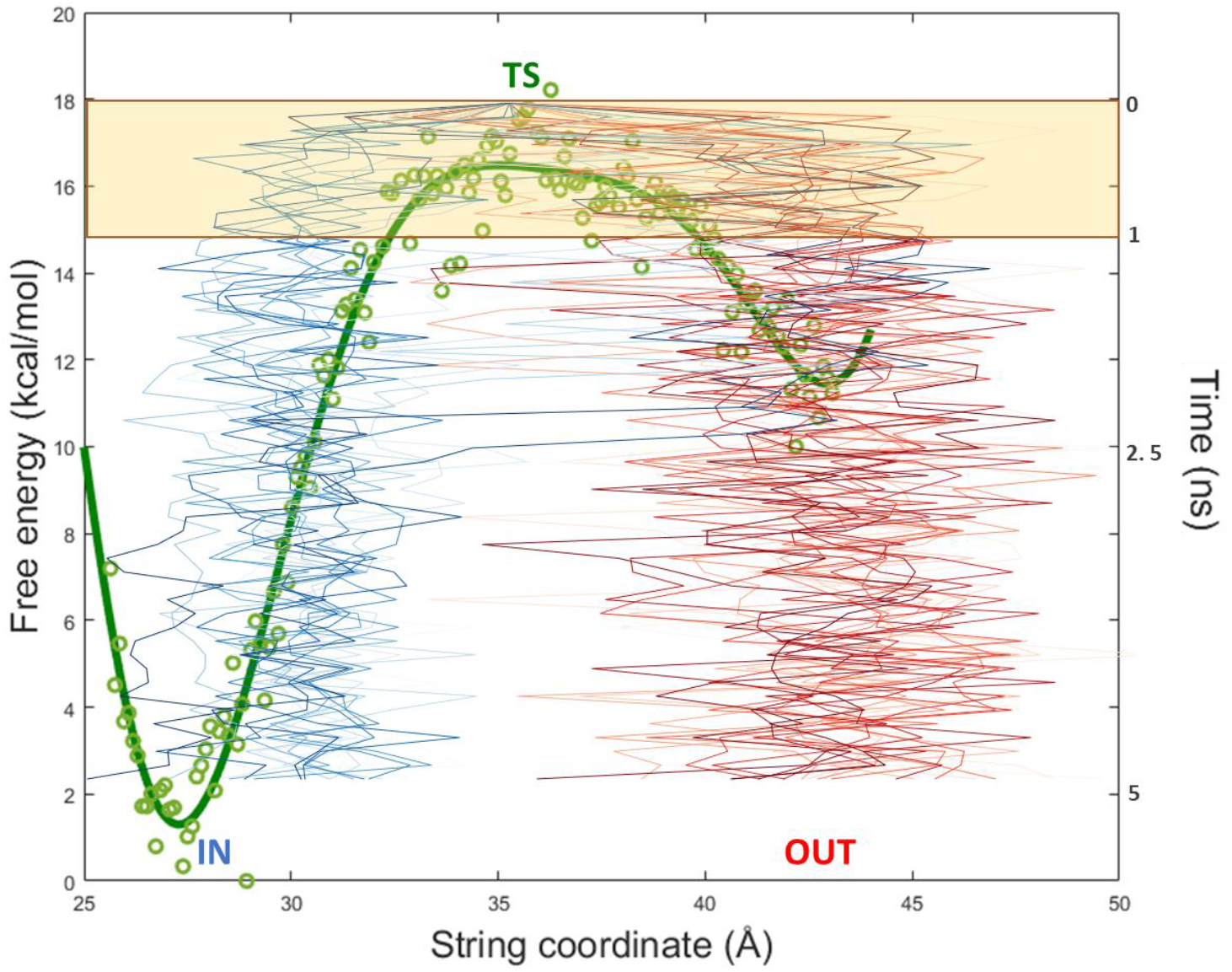
Representation of the PMF of ligand 62K along the string coordinate and the path of multiple downhill trajectories started at the TS (in green) for further analysis. From the TS coordinate as a starting point, a set of simulations leading to both an IN position (blue) and an OUT position (red) are represented as lines. The green dots illustrate the free energy profile datapoints obtained from the WHAM calculation using the string window as string coordinate. The green line represents the fitting obtained from the green dots. The yellow shade represents the simulation time portion used for analysis during our machine learning-based approach.

Using the trained model, we then performed a feature reduction analysis to identify which CV features affect the overall prediction ability of the ML model the most. For both molecules we were able to select the most important structural features (Fig. 8), that lead to the significant reduction of the prediction accuracy, when such features were eliminated (these were kept as a constant value and fed to the ML, see Fig. S3 for details), while other features did not affect the overall accuracy of the predictions.

**Figure 8.**
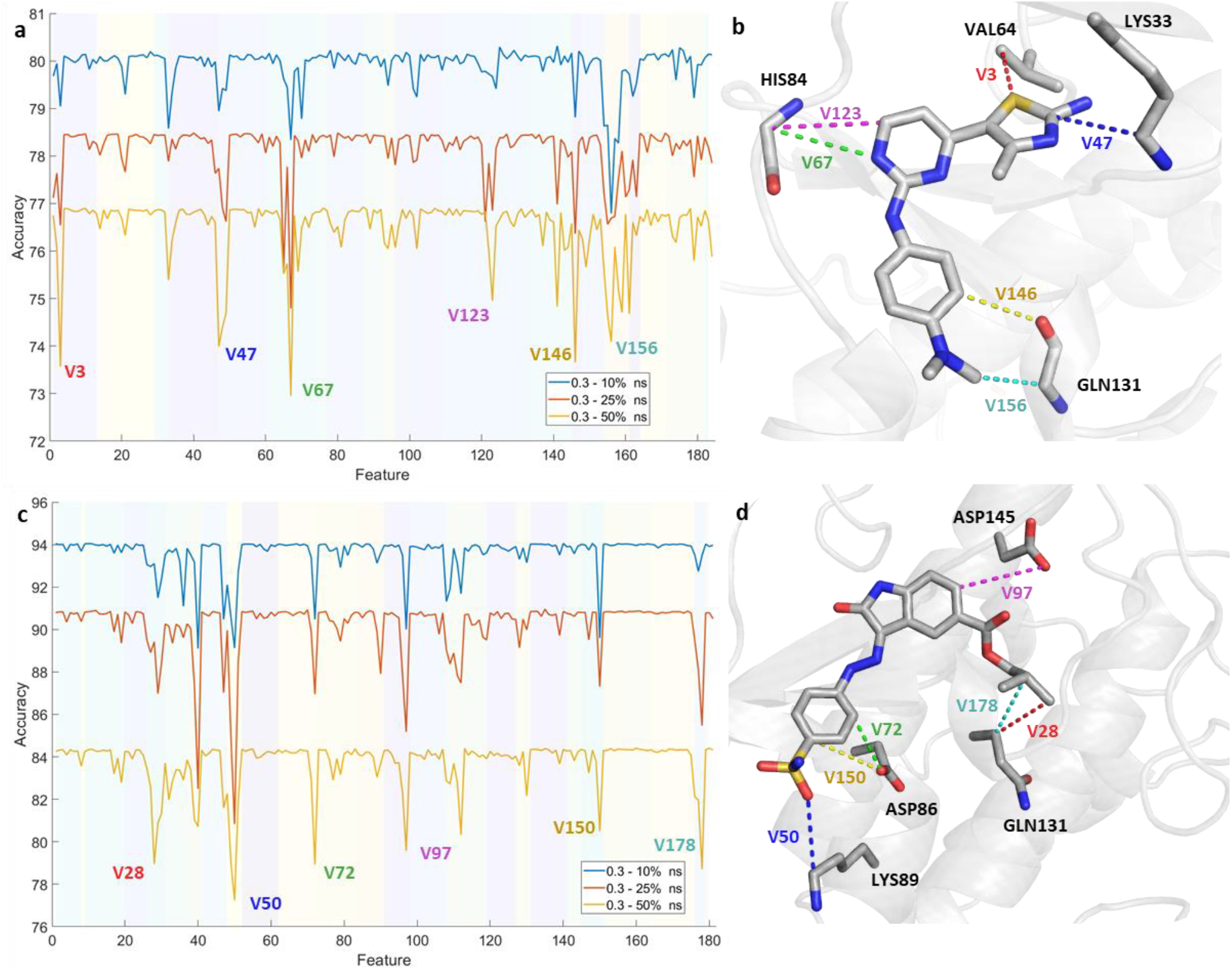
Identification of the essential distances (Feature Reduction) from the largest accuracy drop using the last 50% (yellow), 25% (red), and 10% (blue) of the frames up to the first 0.3 ns of the simulations for **a**: 18K and **c**: 62K. The different shades in the background group the different features according to the atom of the ligand involved. Features presenting significant decrease in accuracy are labelled (see Table S2.II) and portrayed as a 3D representation on the right side of each plot: **b**:18K and **d**:62K.

We also compared the validity of the feature reduction approach with GBDT to identify FIs from the GBDT model. The results obtained show broad similarity with our main MLTSA approach (Figs. S15.I-II) and they both outperform the baseline approach without ML. This suggests that alternative ML models may also be used successfully and further validate our results.

The MLTSA is significantly less computationally intensive than either the unbinding simulations or the string calculations. The short downhill trajectories can be trivially parallelized, which constitute the main cost of the MLTSA analysis. The ML training and accuracy drop calculations have a negligible cost compared to these, therefore MLTSA could be a quick and effective approach to understand key CVs at the TS.

## IV. CONCLUSIONS

Optimizing ligand unbinding kinetics is a very challenging problem for small molecule drug discovery and design, that can lead to the development of drugs with superior efficacy. To tackle this, we have developed a new method, which allows us to calculate the free energy barrier for the ligand unbinding process, therefore providing quantitative information about the residence time of a specific ligand. Our method involves an exploration step, where a ligand unbinding path is determined together with key collective variables that describe this path. Subsequently, we performed accurate free energy calculations using the complete set of identified interactions as CVs along the unbinding path via the finite temperature string method. This provides us with the free energy barriers and an ensemble of structures at the transition state of the ligand unbinding process. The novelty of the method lies in the combination of automated iterative addition and removal of the collective variables determining an unbinding trajectory, which allows us to discover novel interactions not available *a priori*, based on the interactions from the bound structure. We found that while the unbinding trajectories show different paths between different replicas for the same system, our method nevertheless identifies the key interactions important during the unbinding process and provides consistent free energy barriers. The combination of generating an initial path and identifying the important CVs for the unbinding process with the string method for accurate free energy calculations using high dimensional reaction coordinates provide an efficient way to obtain quantitative kinetics of ligand unbinding.

We tested this method using a well-studied cancer drug target, CDK2, using two drug molecules with measured kinetic profiles. We obtained energy barriers in agreement with experiments using our method, which demonstrates the fundamental importance of determining a well-selected, high-dimensional set of CVs for the correct description of the process and kinetics results. We explored analytical 180-dimensional systems using one or multiple DW potentials. We performed the ML analysis both with GBDT and MLP methods. Our results demonstrate for simple linear mixing models that they both can capture correctly the most important correlated features. The MLP is a faster approach and was more sensitive to correlated features, however, sometimes it could not rank the top features in their correct order. On the contrary, the GBDT feature importances could miss lowly correlated features in a dataset but can more accurately rank the top features. The average training time using a single core was around ∼3.5 minutes/model to converge, whereas the GBDT training took about ∼5 minutes/model. Thus, we suggest that a joint approach with both models which may complement one another, could be used to identify relevant CVs. Nonetheless, future studies with non-linear correlated time series can further help to explore the performances of these and other ML methods. Importantly, analogous analysis can be performed for various complex processes, including ones with multiple states as possible outcomes. To aid the kinetics-based design of novel compounds, we also developed a novel method, MLTSA, that allows us to identify the most important features involved at the TS of the unbinding. We generated multiple trajectories initiated at the TS, which either terminated in the bound state or in the unbound state. We then trained a multilayer perceptron ML algorithm to predict the outcome of the trajectories by using a set of CVs and data drawn from the initial segment of the trajectories only. By doing so, we were able to demonstrate that the ML was able to predict the trajectory outcomes with much higher accuracy than using the original set of CVs used for the free energy calculations. A feature importance analysis was further employed to then identify the key CVs and the corresponding structural features that determined the fate of the trajectories, which therefore are the most important descriptors of the TS.

In addition to binding rates, we also aimed to identify specific molecular features and interactions with the target protein that allows us to design kinetic properties of the ligand. Using our ML methods, we identified multiple interactions between the protein and specific parts of the ligands that were of major importance for the trajectories to cross the TS. Important protein-ligand interactions at the TS-bound poses for CDK2 correspond to functional groups of the distal ends of the ligands. Besides His84, a known key residue for interaction with multiple CDK2/4 inhibitors, here we also identified additional common interactions within CDK2 across the ligands, for example between Lys89 and the sulfonamide groups or between Asp145 and the carboxylic group and the ester group for 62K, respectively. Importantly, to perform this analysis, we require the approximate knowledge of the TS structures as well as the MLTSA approach generating a set of downhill unbiased trajectories from these starting points. Our algorithms enable us to uncover novel design objectives for a kinetics-based lead optimization process.

## SUPPLEMENTARY MATERIALS

See Supplemental Material at [URL will be inserted by publisher]. MD simulation details; atom clustering for the unbinding; unbinding CVs; MLTSA flowchart; ML input features; ML training results; free energy profiles; validation of ML analysis; supplementary links to the public GitHub repository containing animated GIF trajectories, code and installable Python package.

## ACKNOWLEDGEMENT

This project made use of time on ARCHER granted via the UK High-End Computing Consortium for Biomolecular Simulation, HECBioSim (http://hecbiosim.ac.uk), supported by EPSRC (grant no. EP/R013012/1) and ERC (project 757850 BioNet).

## For Table of Contents Only

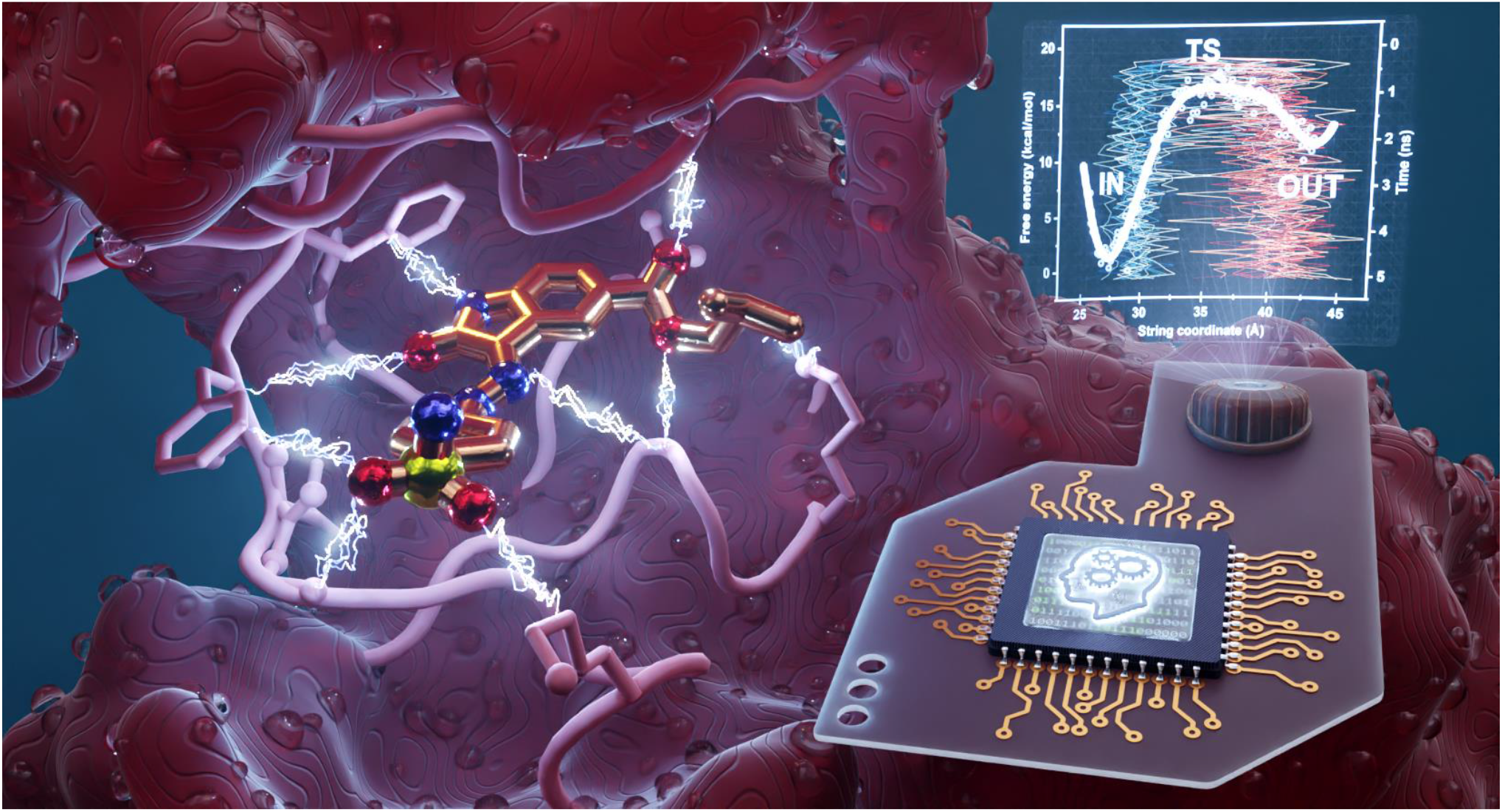

## Supporting Information

### 1. MD Simulations Details

The initial atom coordinates for the three systems were built using high resolution crystal structures with the following PDB codes: **3SW4** (Resolution=1.7 Å), **4FKU** (Resolution=1.47 Å), and **4FKW** (Resolution=1.8 Å)**. We present the results for 4KFU in the SI Section 8.** The systems were modelled using the AMBER ff14SB force field,^1^ and the ligands using the general Amber force field (GAFF).^2^ The ligand’s atomic partial charges were obtained using density functional theory (DFT) ωB97X-D/def2TZVPP^3,4^ as implemented in Gaussian 09 Revision E.^5^ The full system was solvated with 12,000-14,000 TIP3P water molecules. Na^+^ and Cl^-^ ions were added to neutralize the system and set a salt concentration of 0.14 M. All the MD simulations were performed using NAMD 2.12.^6^

The three systems were first minimized using a standard protocol via the steepest descent algorithm for a total of 150,000 steps followed by the equilibration of the restrained protein (1 kcal mol^-1^ Å^-2^ force applied to each heavy atom of the protein) for 10 ns in NVT ensemble at 300 K via a standard MD procedure. All the production runs were performed with the NPT ensemble with a time step of 2 fs. Pressure was maintained at 1 atm by a Nosé–Hoover Langevin piston.^7^ Temperature was maintained at 298 K using Langevin dynamics with a damping coefficient *γ* of 0.5 ps^-1^ applied to all atoms. SHAKE^8^ was applied to all bonds involving hydrogen and nonbonded interactions were calculated with a cutoff of 12 Å, and a switching distance of 10 Å. The particle mesh Ewald method was used for long-range electrostatic calculations with a grid density of >1 Å^−3.9^

An initial unbiased simulation of 20 ns was performed for each ligand. This initial simulation allows the system to equilibrate and gives us an initial trajectory to calculate the first CVs.

### 2. Atom Clustering

Residues with atoms that have a rotational degree of freedom with multiple equivalent positions are clustered together. During the unbinding process, if a new contact is found with one atom belonging to the cluster, then the harmonic restraint will be applied to the centre of mass of the selected clustered atoms. The use of clustered atoms reduces the fluctuation caused by the rotation of such bonds, affecting the overall harmonic restraint.

**Figure S1.**
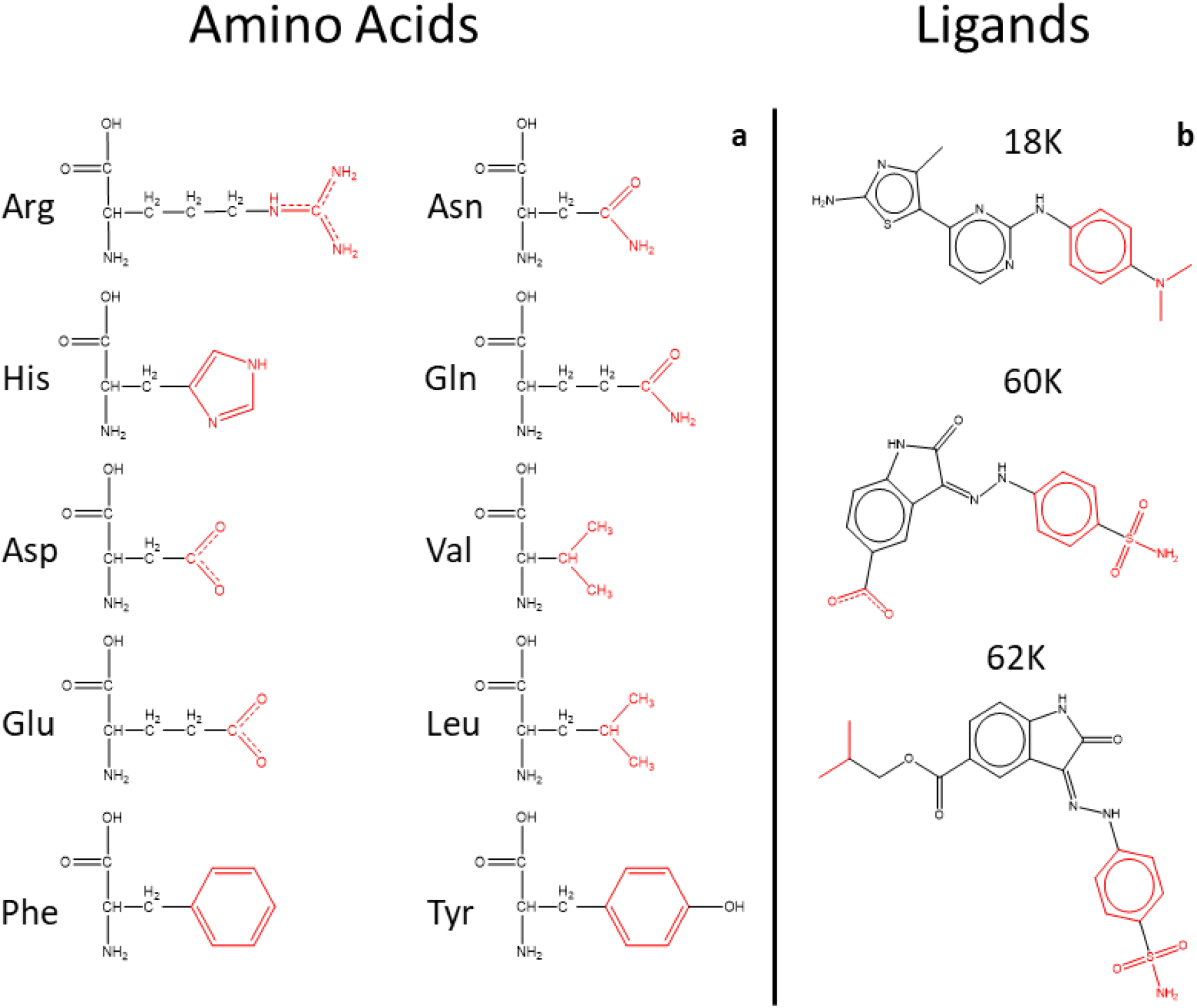
Chemical structures of the residues with clustered atoms, highlighted in red; a for the amino acids and b for the ligands.

### 3. Unbinding Collective Variables

Distances are included and excluded throughout the entire unbinding trajectory. The initial distances are calculated from the initial unbiased ligand-bound MD simulations. At every 10 ns, the biased simulation is stopped and analyzed. New close interactions between the protein and ligand will be added as part of the main CV that is biased, and previous CVs where the values exceeded the cut-off are excluded. Fig. S2.I-S2.VI display the distances that are part of the main CV along the unbinding trajectories for each ligand and each replica.

**Figure S2.I.**
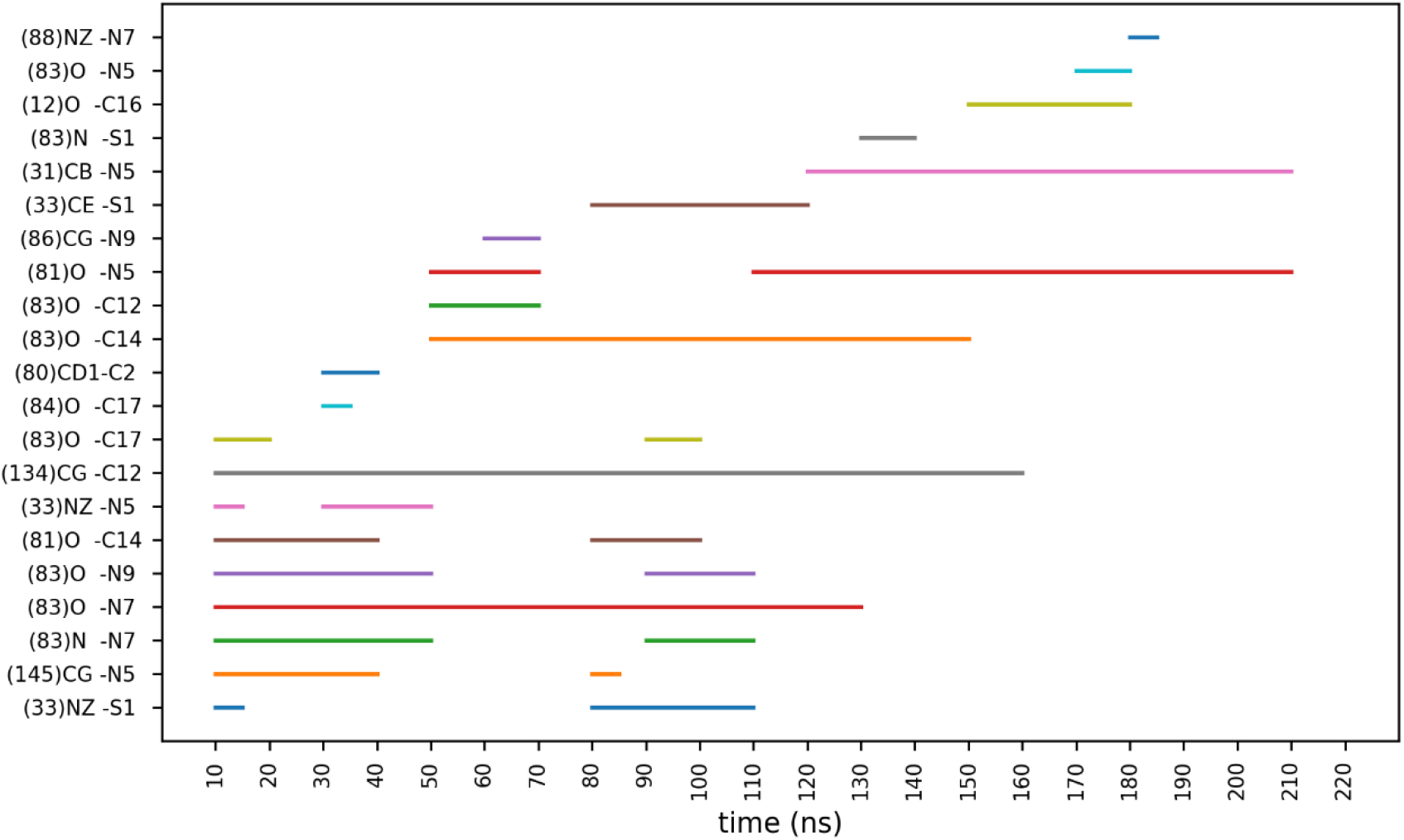
Distances used for the main CV along the trajectory for 18K replica 1.

**Figure S2.II.**
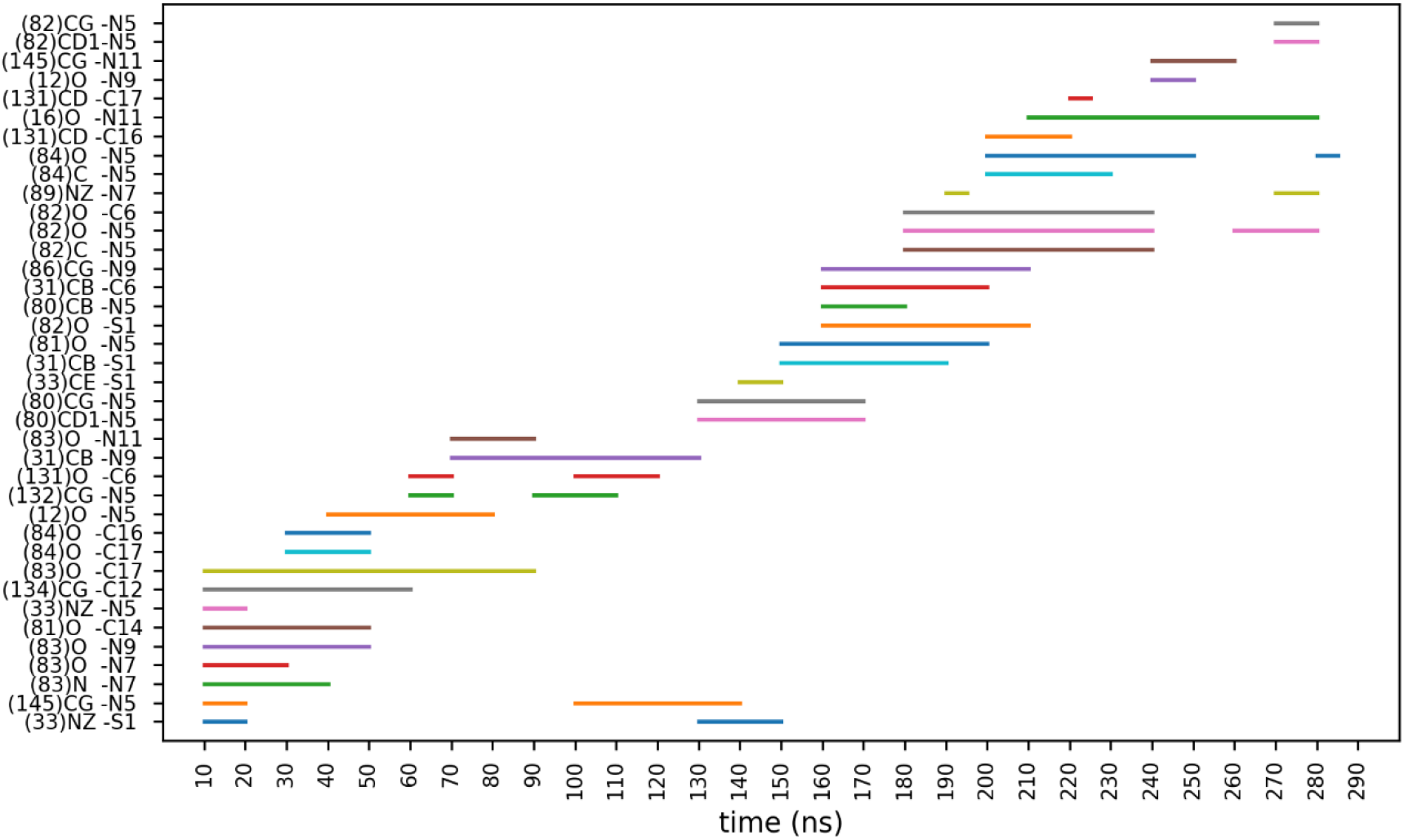
Distances used for the main CV along the trajectory for 18K replica 2.

**Figure S2.III.**
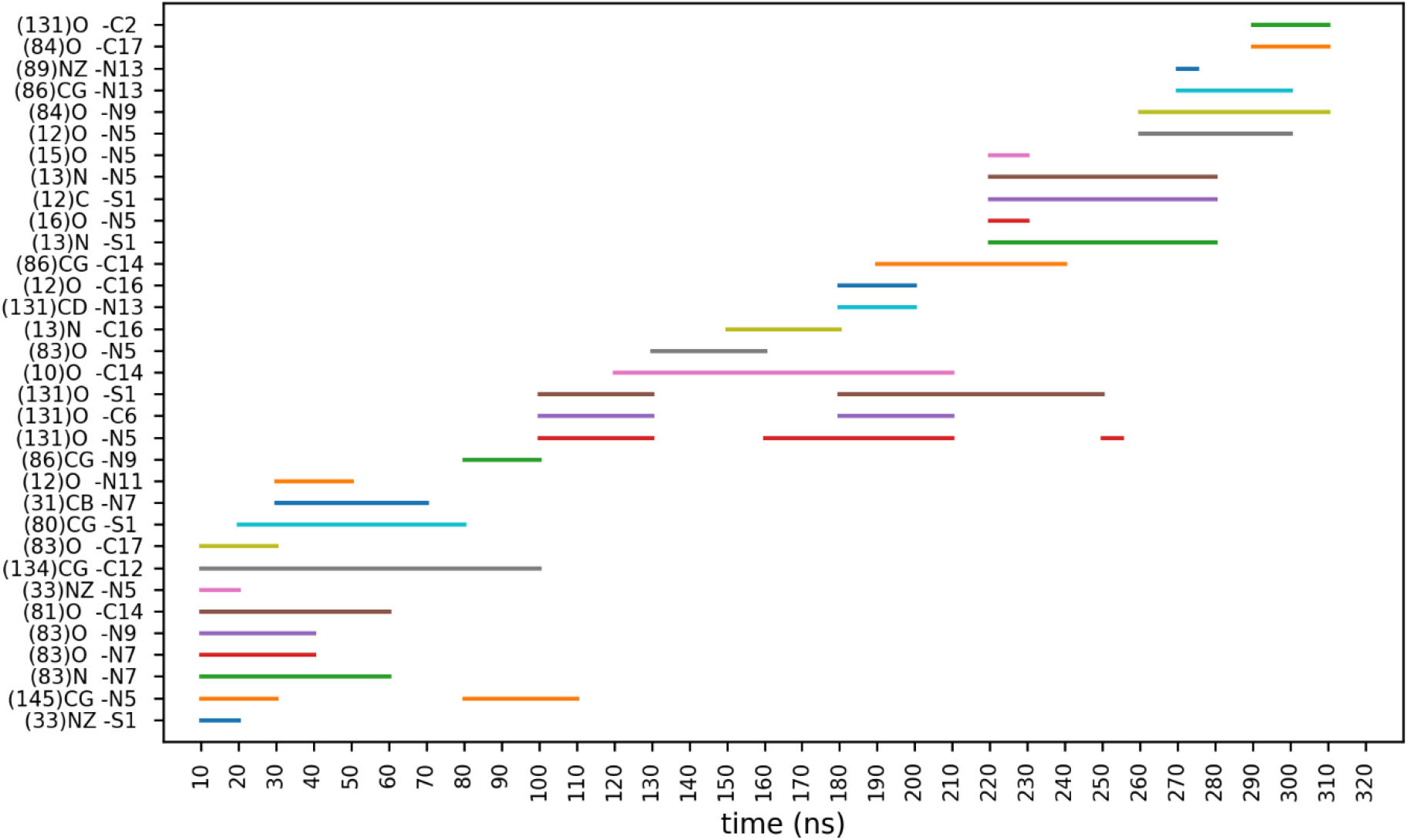
Distances used for the main CV along the trajectory 18K replica 3.

**Figure S2.IV.**
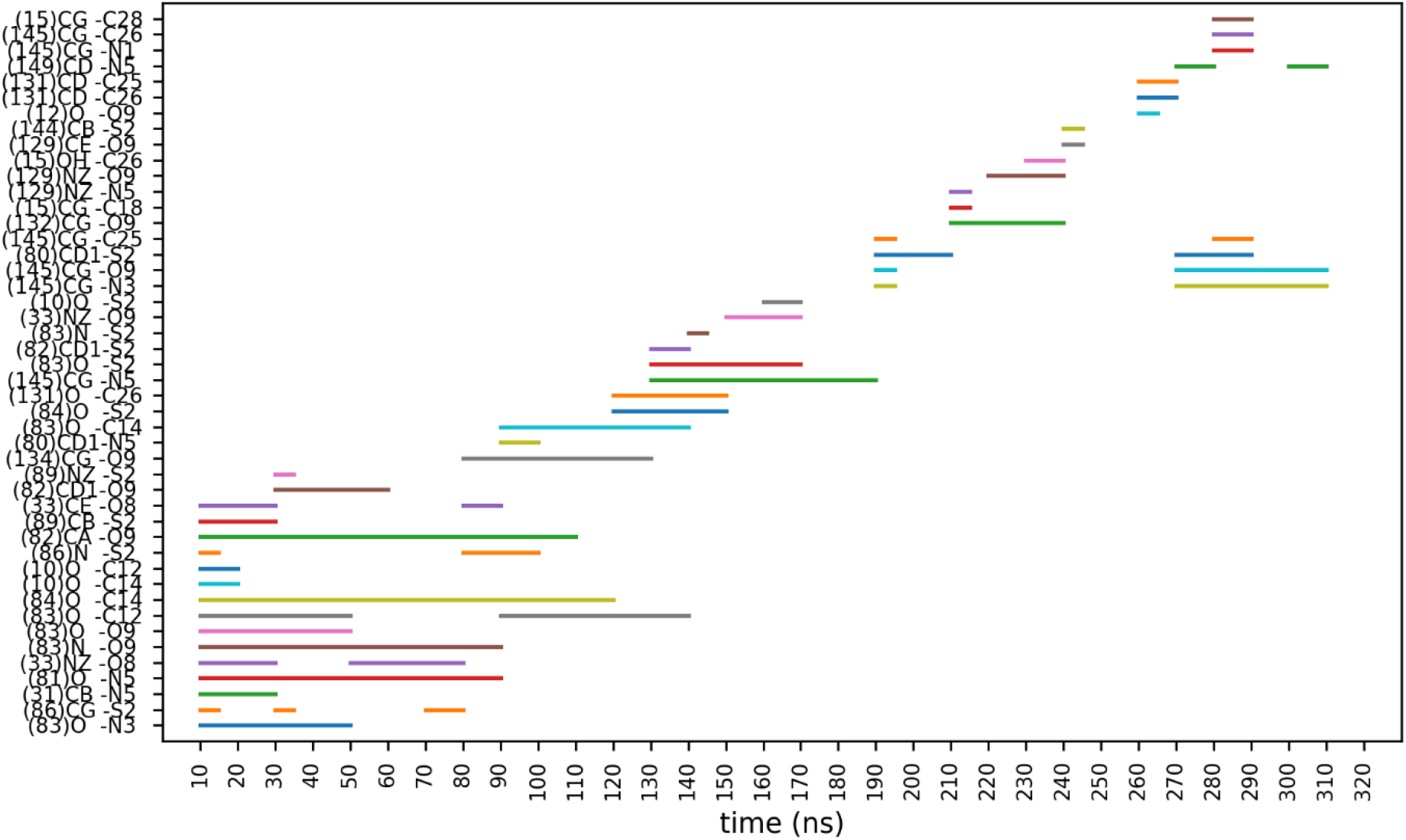
Distances used for the main CV along the trajectory for 62K replica 1.

**Figure S2.V.**
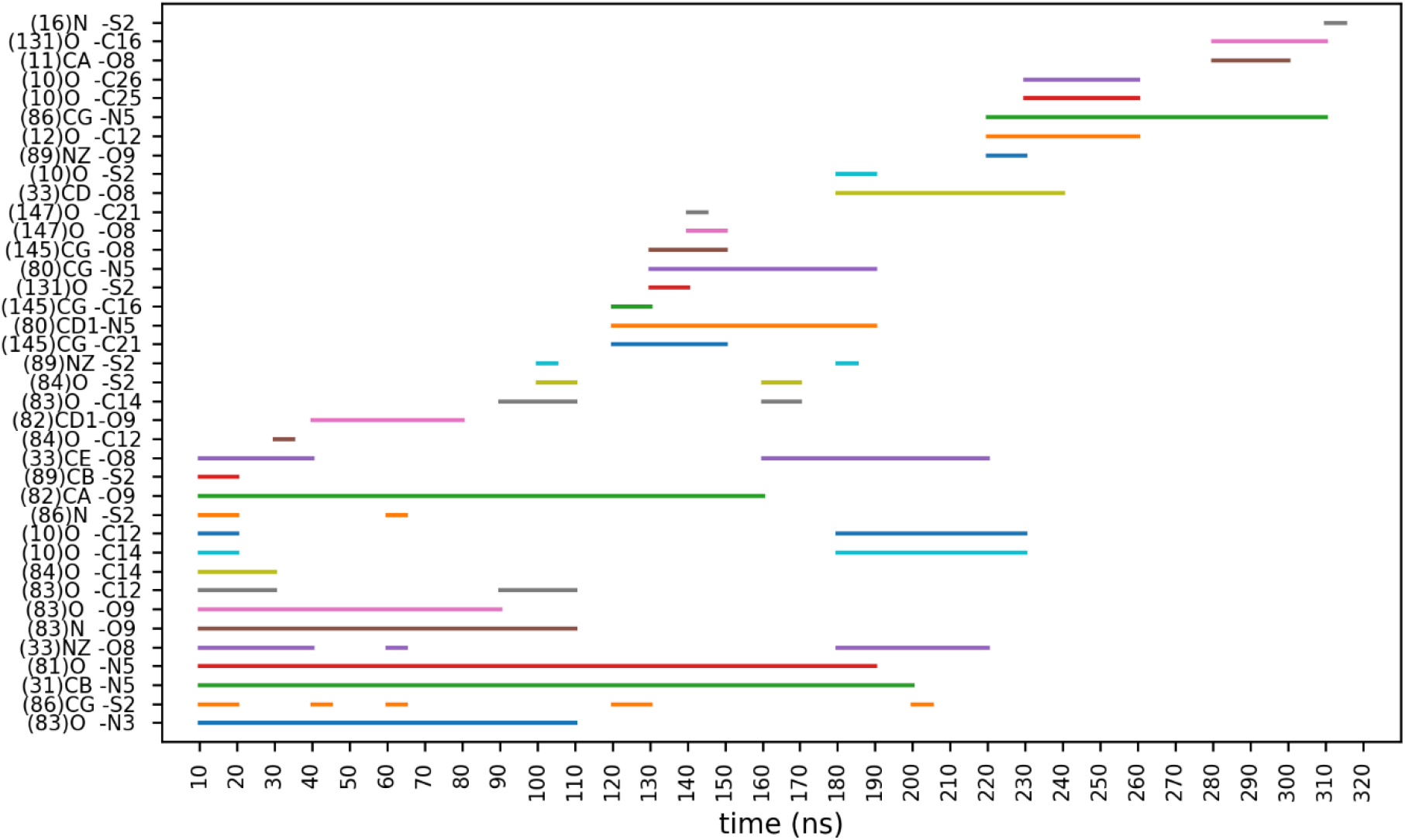
Distances used for the main CV along the trajectory for 62K replica 2.

**Figure S2.VI.**
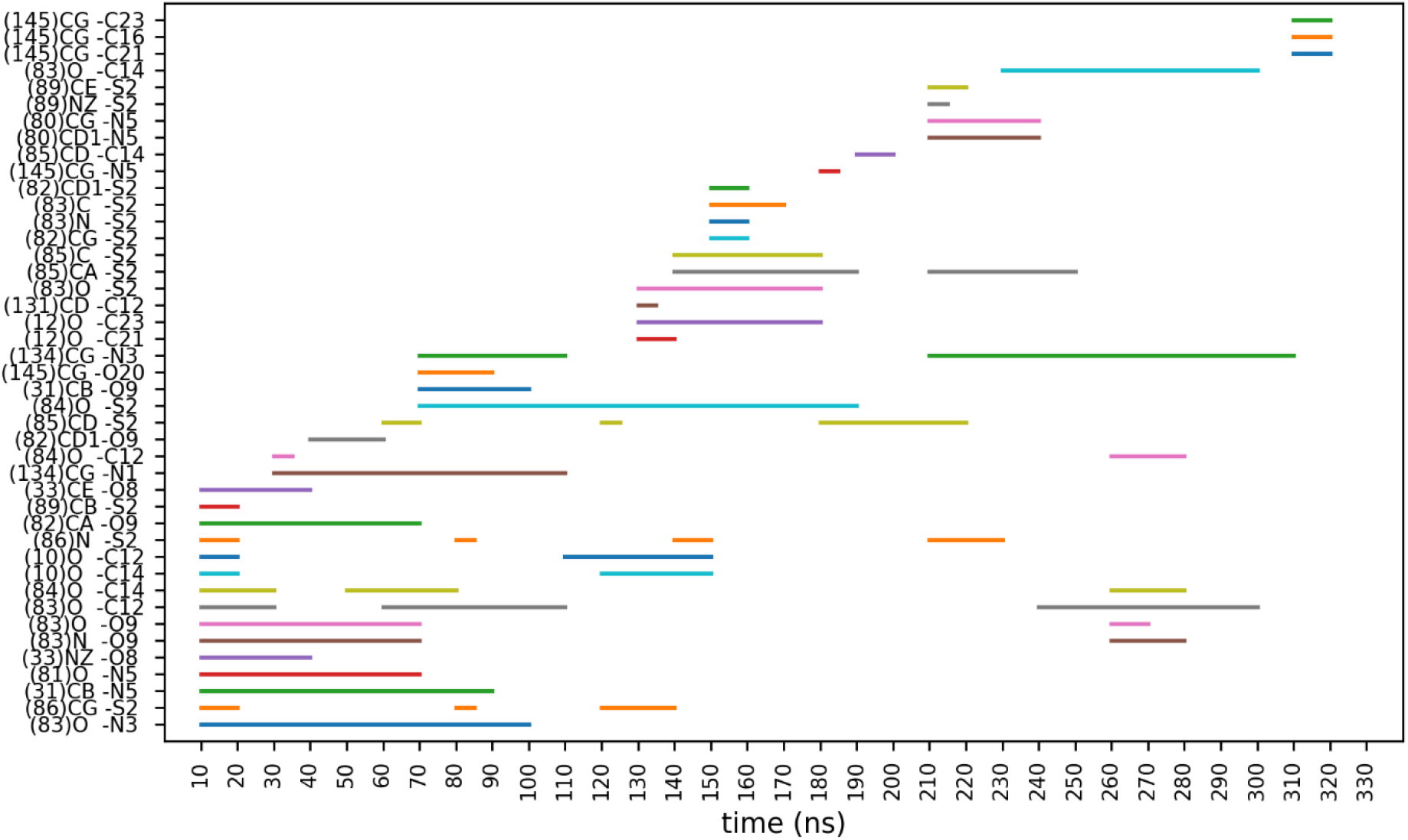
Distances used for the main CV along the trajectory for 62K replica 3.

### 4. MLTSA flowchart

**Figure S3.**
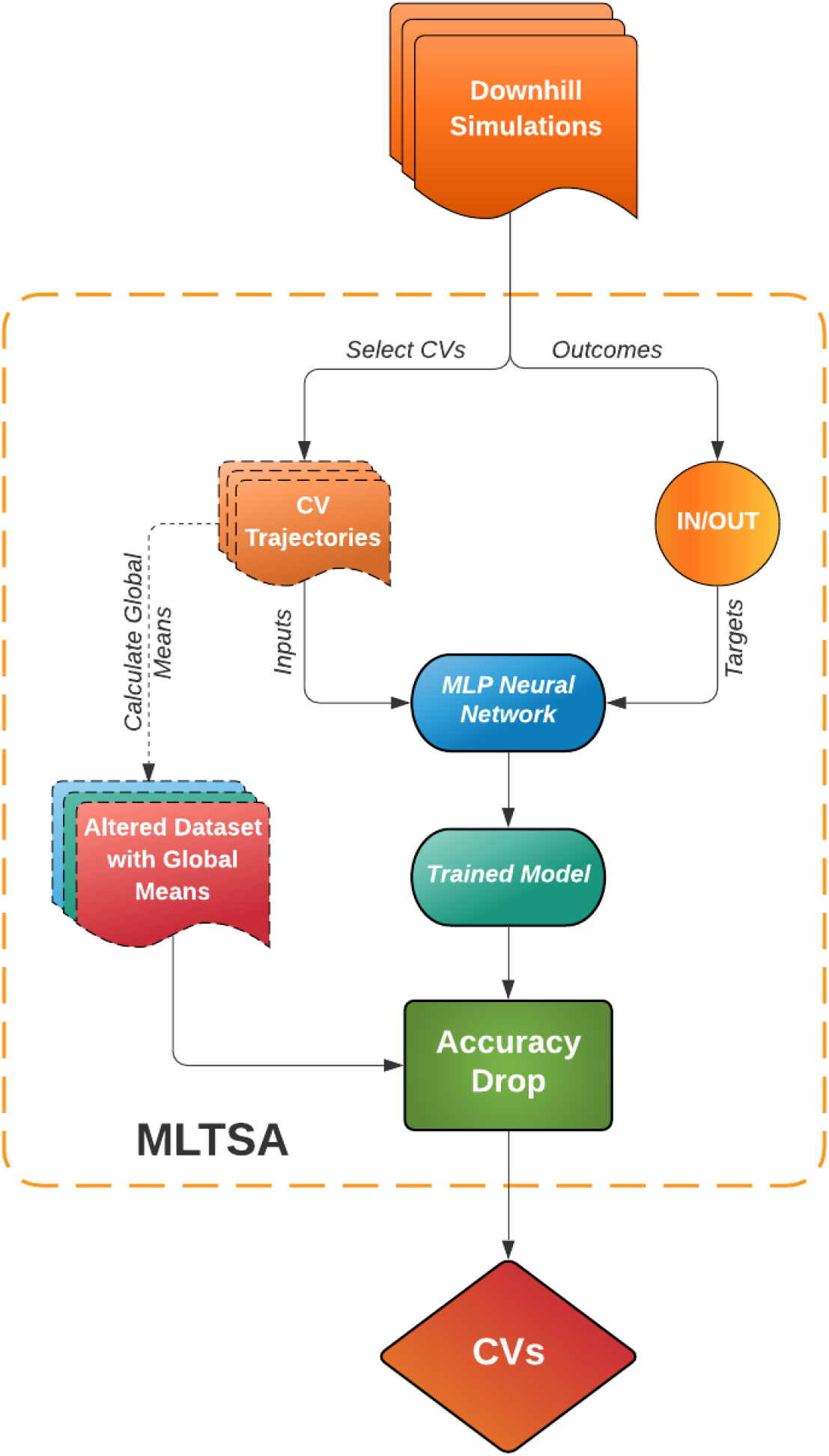
Flowchart of the Machine Learning Transition State Analysis (MLTSA) process. This approach uses data from the downhill unbiased simulations initiated near the TS to train a neural network on the trajectory frames using selected CVs. The model then predicts the ligand’s outcome (In/Out). After successful training, the original input data (orange) is altered one CV at a time and replaced by a constant value (Global Mean), thus producing an altered dataset (Red/Green/Blue). This is used to re-predict the outcomes on the already trained model (Sea Green) and pinpoint the main accuracy drops. This allows identifying the most relevant CVs by assessing the accuracy drop.

### 5. Classification and Input Features for Machine Learning

#### Analytical model system

Two types of 1D potentials were used: single-well (SW, *y_sw_*) potentials and double-well (DW, *y_dw_*) potentials, defined by the equation:

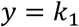

where *k*_1_ = 100, μ = ½, *σ* = 0.01, and *k*_2_= 0 for a SW potential or **k**_2_= 1 for a DW potential.

To generate trajectories along these model 1D free energy profiles, we used the 1D Brownian Overdamped Langevin Equation:

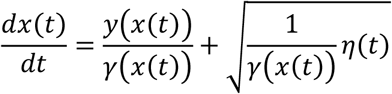

where *γ* = 0.01is constant along *x* and *η*(*t*) is a number randomly sampled from a normal (Gaussian) distribution centred at 0 and the spread is 1.0.

Using the trajectories generated, we defined input features (*y_feature_*) by combining the coordinates along two different potentials (*y_pot_*), either SW or DW using *y_feature_* = **±*y_pot1_* + (1 − *±*)*y_pot2_*. We generated 180 features for both a dataset with 24 SW potentials and 1 single DW potential which decides the IN/OUT state, and for a dataset with 20 SW potentials and 5 DW potentials, having among them the decisive DW potential as well.

**Table S1.I.** Table containing the feature numbers for the correlated features, the values of the mixing coefficients (**α_1_**) for the selected DW potential, and the second SW potential used for the linear combination. Coefficients for all other uncorrelated input features are not shown.

| Feature Number | $\alpha_1$ | SW |
| --- | --- | --- |
| 5 | **0.575** | <u>11</u> |
| 21 | **0.475** | <u>10</u> |
| 25 | **0.535** | <u>21</u> |
| 42 | **0.673** | <u>19</u> |
| 48 | **0.403** | <u>23</u> |
| 55 | **0.840** | <u>18</u> |
| 87 | **0.656** | <u>1</u> |
| 90 | **0.348** | <u>4</u> |
| 119 | **0.858** | <u>12</u> |
| 136 | **0.355** | <u>9</u> |
| 167 | **0.899** | <u>3</u> |

**Table S1.II.**
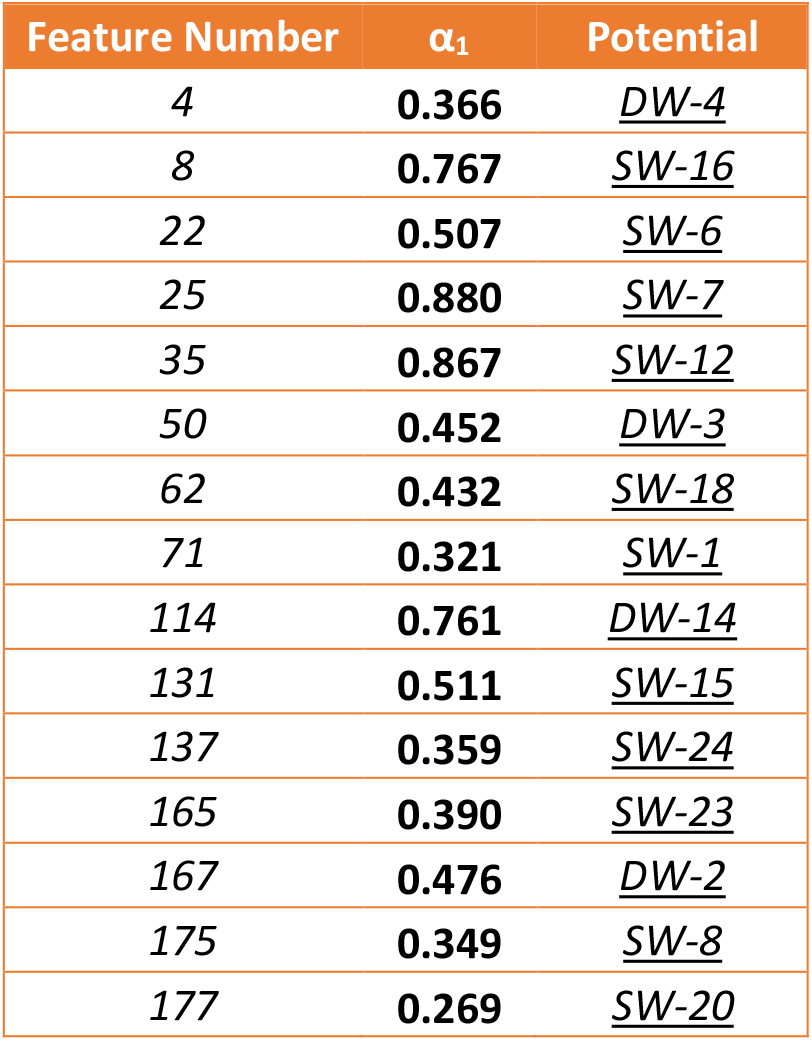
Table containing the feature numbers for the correlated features, the values of the mixing coefficients (**α_1_**) for the selected DW potential, and the second SW or one of the five DW potentials used for the linear combination. Coefficients for all other uncorrelated input features are not shown.

**Figure S4.**
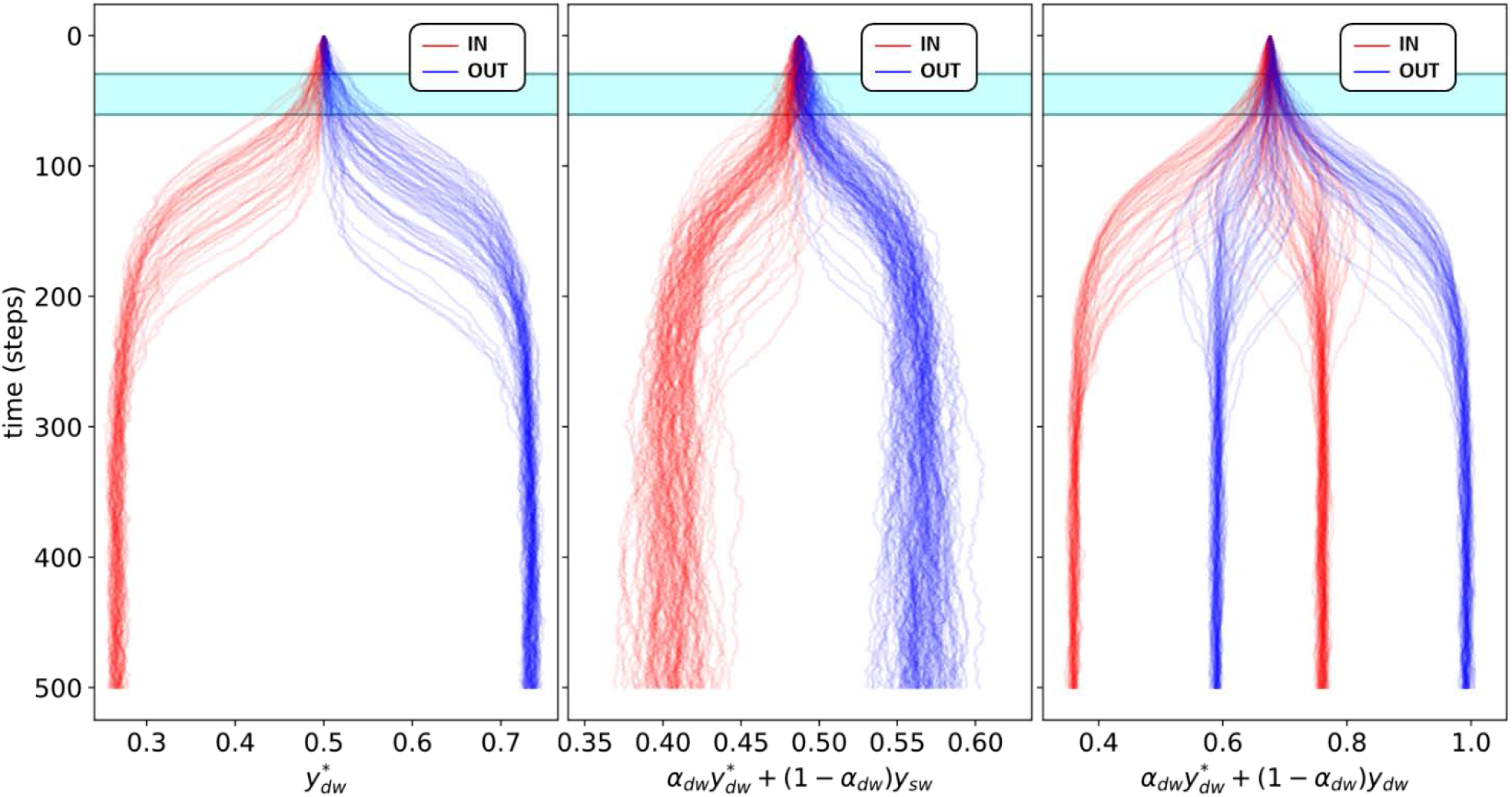
Left: Plot of the trajectory coordinates across different simulations depending on the classified outcome (IN-red OUT-blue) for the decisive DW potential. Middle: Plot of one correlated feature for different simulations after the linear combination between the decisive DW and a SW potential, this plot corresponds to feature 136 of the 1DW dataset, with a linear coefficient of α = 0.355. Right: Plot of a correlated feature which is combined with the decisive DW potential and another DW potential, the line colour depends on the classified outcome. In this case this corresponds to feature 4 from the 5 DW dataset, the decisive DW (marked with *) has a linear coefficient of α = 0.366. In shaded cyan the region selected for the ML training (from the 30th step to the 60th).

#### CDK2 system data pre-processing

The classification of the ligand in the bound position (IN) and unbound position (OUT) is calculated by analyzing the last 250 ps of the downhill trajectories. For each frame we extract and sum two key distances between the ligand and the protein (see Table S2.I) and average these for all the frames of the last 250 ps. If the sum of these distances is below a given IN-threshold the trajectory is classified as IN, if the value is above the OUT-threshold then is classified as OUT (see Table S2.I). Table S2.II presents all the interatomic distances used in the ML inputs.

**Table S2.I.** Key distances used to automate the IN/OUT labelling of the 5 ns-long downhill trajectories. These are used to create a dataset suitable for the ML algorithm to learn the classification with the selected CVs as inputs (X) and the labels IN/OUT as targets (Y).

| SYSTEM | DISTANCES | IN-THRESHOLD | OUT-THRESHOLD |
| --- | --- | --- | --- |
| 3SW4 | LIG(N9)-LEU83(O) -<br>LIG(N7)-LEU83(N) | 10 | 12 |
| 4FKW | LIG(N3)-LEU83(O) -<br>LIG(O9)-LEU83(N) | 11 | 13 |

**Table S2.II.** List of all interatomic distances for heavy atoms between the ligand and the protein within 6 Å at the selected TS structure. In addition, three intra-residue distances within the protein were also added.

| **3sw4**<br>**(184 Features)** | **4fkw**<br>**(181 Features)** |
| --- | --- |
| ===== | ===== |
| V1=LIG(S1)-VAL64(CB) | V1=LIG(N1)-ILE10(CB) |
| V2=LIG(S1)-VAL64(CG1) | V2=LIG(N1)-ILE10(CG1) |
| V3=LIG(S1)-VAL64(CG2) | V3=LIG(N1)-ILE10(CD1) |
| V4=LIG(S1)-PHE80(CB) | V4=LIG(N1)-GLN131(O) |
| V5=LIG(S1)-PHE80(CG) | V5=LIG(N1)-LEU134(CG) |
| V6=LIG(S1)-PHE80(CD1) | V6=LIG(N1)-LEU134(CD1) |
| V7=LIG(S1)-PHE80(CE1) | V7=LIG(N1)-LEU134(CD2) |
| V8=LIG(S1)-PHE80(CZ) | V8=LIG(S2)-ILE10(CG2) |
| V9=LIG(S1)-PHE80(CE2) | V9=LIG(N3)-ILE10(CB) |
| V10=LIG(S1)-PHE80(CD2) | V10=LIG(N3)-ILE10(CG1) |
| V11=LIG(S1)-LEU134(CD1) | V11=LIG(N3)-ILE10(CD1) |
| V12=LIG(S1)-ALA144(CB) | V12=LIG(N3)-GLN85(O) |
| V13=LIG(C2)-ILE10(CB) | V13=LIG(N3)-ASP86(CB) |
| V14=LIG(C2)-ILE10(CG2) | V14=LIG(N3)-LEU134(CG) |
| V15=LIG(C2)-ILE10(CD1) | V15=LIG(N3)-LEU134(CD1) |
| V16=LIG(C2)-VAL18(CB) | V16=LIG(N3)-LEU134(CD2) |
| V17=LIG(C2)-VAL18(CG1) | V17=LIG(N4)-ILE10(CA) |
| V18=LIG(C2)-VAL18(CG2) | V18=LIG(N4)-ILE10(CB) |
| V19=LIG(N3)-VAL18(CG2) | V19=LIG(N4)-ILE10(CG2) |
| V20=LIG(N3)-LYS33(CE) | V20=LIG(N5)-ILE10(CD1) |
| V21=LIG(N3)-LYS33(NZ) | V21=LIG(N5)-VAL18(CG2) |
| V22=LIG(N3)-PHE80(CG) | V22=LIG(N5)-ALA31(CB) |
| V23=LIG(N3)-PHE80(CD1) | V23=LIG(N5)-LEU134(CD1) |
| V24=LIG(N3)-PHE80(CE1) | V24=LIG(N5)-LEU134(CD2) |
| V25=LIG(N3)-PHE80(CZ) | V25=LIG(N5)-ALA144(CB) |
| V26=LIG(N3)-LEU134(CD1) | V26=LIG(C6)-LYS129(CD) |
| V27=LIG(N3)-ALA144(CB) | V27=LIG(C6)-LYS129(CE) |
| V28=LIG(N3)-ASP145(OD2) | V28=LIG(C6)-LYS129(NZ) |
| V29=LIG(C4)-VAL18(CB) | V29=LIG(C6)-GLN131(CB) |
| V30=LIG(C4)-VAL18(CG1) | V30=LIG(C6)-GLN131(CG) |
| V31=LIG(C4)-VAL18(CG2) | V31=LIG(C7)-ILE10(O) |
| V32=LIG(C4)-LEU134(CD1) | V32=LIG(C7)-GLY11(N) |
| V33=LIG(N5)-LYS33(CE) | V33=LIG(C7)-GLY11(CA) |
| V34=LIG(N5)-LYS33(NZ) | V34=LIG(C7)-GLY11(C) |
| V35=LIG(N5)-VAL64(CG1) | V35=LIG(C7)-GLU12(N) |
| V36=LIG(N5)-PHE80(CB) | V36=LIG(C7)-GLN131(CB) |
| V37=LIG(N5)-PHE80(CG) | V37=LIG(O8)-VAL18(CG1) |
| V38=LIG(N5)-PHE80(CD1) | V38=LIG(O8)-ASP145(CB) |
| V39=LIG(N5)-PHE80(CE1) | V39=LIG(O8)-ASP145(CG) |
| V40=LIG(N5)-PHE80(CZ) | V40=LIG(O8)-ASP145(OD2) |
| V41=LIG(N5)-PHE80(CE2) | V41=LIG(O9)-ILE10(CD1) |
| V42=LIG(N5)-PHE80(CD2) | V42=LIG(O9)-ALA31(CB) |
| V43=LIG(N5)-ALA144(CA) | V43=LIG(O9)-GLU81(O) |
| V44=LIG(N5)-ALA144(CB) | V44=LIG(O9)-LEU134(CG) |
| V45=LIG(N5)-ASP145(N) | V45=LIG(O9)-LEU134(CD1) |
| V46=LIG(N5)-ASP145(OD2) | V46=LIG(O9)-LEU134(CD2) |
| V47=LIG(C6)-LYS33(CE) | V47=LIG(O10)-HIE84(O) |
| V48=LIG(C6)-VAL64(CG1) | V48=LIG(O11)-ASP86(OD2) |
| V49=LIG(C6)-VAL64(CG2) | V49=LIG(O11)-LYS89(CG) |
| V50=LIG(C6)-PHE80(CB) | V50=LIG(O11)-LYS89(CD) |
| V51=LIG(C6)-PHE80(CG) | V51=LIG(O11)-LYS89(CE) |
| V52=LIG(C6)-PHE80(CD1) | V52=LIG(C12)-ILE10(CB) |
| V53=LIG(C6)-PHE80(CE1) | V53=LIG(C12)-ILE10(CG2) |
| V54=LIG(C6)-PHE80(CZ) | V54=LIG(C12)-ILE10(CG1) |
| V55=LIG(C6)-PHE80(CE2) | V55=LIG(C12)-ILE10(CD1) |
| V56=LIG(C6)-PHE80(CD2) | V56=LIG(C12)-PHE82(CE1) |
| V57=LIG(C6)-LEU134(CD1) | V57=LIG(C12)-LEU83(O) |
| V58=LIG(C6)-ALA144(CA) | V58=LIG(C12)-GLN85(N) |
| V59=LIG(C6)-ALA144(CB) | V59=LIG(C12)-GLN85(CA) |
| V60=LIG(N7)-ILE10(CG2) | V60=LIG(C12)-GLN85(C) |
| V61=LIG(N7)-ILE10(CD1) | V61=LIG(C12)-GLN85(O) |
| V62=LIG(N7)-LEU83(N) | V62=LIG(C13)-ILE10(CB) |
| V63=LIG(N7)-LEU83(CA) | V63=LIG(C13)-ILE10(CG2) |
| V64=LIG(N7)-LEU83(CB) | V64=LIG(C13)-ILE10(CG1) |
| V65=LIG(N7)-LEU83(C) | V65=LIG(C13)-ILE10(CD1) |
| V66=LIG(N7)-HIE84(N) | V66=LIG(C13)-ILE10(O) |
| V67=LIG(N7)-HIE84(CA) | V67=LIG(C13)-GLN85(C) |
| V68=LIG(N7)-HIE84(C) | V68=LIG(C13)-ASP86(N) |
| V69=LIG(N7)-HIE84(O) | V69=LIG(C13)-ASP86(CA) |
| V70=LIG(N7)-GLN85(N) | V70=LIG(C13)-ASP86(CB) |
| V71=LIG(N7)-GLN85(C) | V71=LIG(C13)-ASP86(CG) |
| V72=LIG(N7)-GLN85(O) | V72=LIG(C13)-ASP86(OD2) |
| V73=LIG(N7)-LEU134(CD1) | V73=LIG(C14)-ILE10(CB) |
| V74=LIG(N7)-LEU134(CD2) | V74=LIG(C14)-ILE10(CG2) |
| V75=LIG(C8)-ALA31(CB) | V75=LIG(C14)-ILE10(CD1) |
| V76=LIG(C8)-LEU134(CD1) | V76=LIG(C14)-PHE82(CE1) |
| V77=LIG(N9)-ILE10(CG2) | V77=LIG(C14)-LEU83(O) |
| V78=LIG(N9)-HIE84(C) | V78=LIG(C14)-HIE84(C) |
| V79=LIG(N9)-HIE84(O) | V79=LIG(C14)-HIE84(O) |
| V80=LIG(N9)-GLN85(C) | V80=LIG(C14)-GLN85(N) |
| V81=LIG(N9)-GLN85(O) | V81=LIG(C14)-GLN85(CA) |
| V82=LIG(N9)-ASP86(N) | V82=LIG(C14)-GLN85(C) |
| V83=LIG(N9)-ASP86(CA) | V83=LIG(C15)-ILE10(CB) |
| V84=LIG(N9)-LEU134(CG) | V84=LIG(C15)-ILE10(CG2) |
| V85=LIG(N9)-LEU134(CD1) | V85=LIG(C15)-ILE10(O) |
| V86=LIG(N9)-LEU134(CD2) | V86=LIG(C15)-GLN85(C) |
| V87=LIG(C10)-ILE10(CG2) | V87=LIG(C15)-ASP86(N) |
| V88=LIG(C10)-ILE10(CD1) | V88=LIG(C15)-ASP86(CB) |
| V89=LIG(C10)-ALA31(CB) | V89=LIG(C15)-ASP86(CG) |
| V90=LIG(C10)-LEU134(CG) | V90=LIG(C15)-ASP86(OD2) |
| V91=LIG(C10)-LEU134(CD1) | V91=LIG(C16)-VAL18(CB) |
| V92=LIG(C10)-LEU134(CD2) | V92=LIG(C16)-VAL18(CG1) |
| V93=LIG(N11)-ASP86(CG) | V93=LIG(C16)-VAL18(CG2) |
| V94=LIG(N11)-ASP86(OD1) | V94=LIG(C16)-ALA144(CB) |
| V95=LIG(N11)-ASP86(OD2) | V95=LIG(C16)-ASP145(CB) |
| V96=LIG(C12)-ILE10(CD1) | V96=LIG(C16)-ASP145(CG) |
| V97=LIG(C12)-ALA31(CB) | V97=LIG(C16)-ASP145(OD2) |
| V98=LIG(C12)-GLU81(O) | V98=LIG(C17)-VAL18(CB) |
| V99=LIG(C12)-LEU83(N) | V99=LIG(C17)-VAL18(CG1) |
| V100=LIG(C12)-LEU83(CA) | V100=LIG(C17)-VAL18(CG2) |
| V101=LIG(C12)-LEU83(CB) | V101=LIG(C17)-ALA144(CB) |
| V102=LIG(C12)-LEU83(CG) | V102=LIG(C17)-ASP145(CB) |
| V103=LIG(C12)-LEU134(CD1) | V103=LIG(C18)-VAL18(CB) |
| V104=LIG(N13)-ILE10(CG2) | V104=LIG(C18)-VAL18(CG1) |
| V105=LIG(N13)-ILE10(CD1) | V105=LIG(C18)-VAL18(CG2) |
| V106=LIG(N13)-LEU134(CG) | V106=LIG(C18)-GLN131(O) |
| V107=LIG(N13)-LEU134(CD1) | V107=LIG(C18)-LEU134(CD2) |
| V108=LIG(N13)-LEU134(CD2) | V108=LIG(C19)-LYS129(CE) |
| V109=LIG(C14)-ILE10(CG2) | V109=LIG(C19)-LYS129(NZ) |
| V110=LIG(C14)-ILE10(CD1) | V110=LIG(C19)-GLN131(CA) |
| V111=LIG(C14)-ALA31(CB) | V111=LIG(C19)-GLN131(CB) |
| V112=LIG(C14)-GLU81(O) | V112=LIG(C19)-GLN131(CG) |
| V113=LIG(C14)-PHE82(CA) | V113=LIG(C19)-GLN131(C) |
| V114=LIG(C14)-PHE82(CD1) | V114=LIG(C19)-GLN131(O) |
| V115=LIG(C14)-PHE82(CE1) | V115=LIG(C19)-ASN132(N) |
| V116=LIG(C14)-PHE82(C) | V116=LIG(C19)-ASN132(CG) |
| V117=LIG(C14)-LEU83(N) | V117=LIG(C19)-ASN132(OD1) |
| V118=LIG(C14)-LEU83(CA) | V118=LIG(C19)-ASN132(ND2) |
| V119=LIG(C14)-LEU83(CB) | V119=LIG(O20)-GLN131(CB) |
| V120=LIG(C14)-LEU83(CG) | V120=LIG(O20)-GLN131(C) |
| V121=LIG(C14)-LEU83(C) | V121=LIG(O20)-GLN131(O) |
| V122=LIG(C14)-HIE84(N) | V122=LIG(O20)-ASN132(N) |
| V123=LIG(C14)-HIE84(CA) | V123=LIG(O20)-ASN132(CA) |
| V124=LIG(C14)-HIE84(C) | V124=LIG(O20)-ASN132(CG) |
| V125=LIG(C14)-HIE84(O) | V125=LIG(O20)-ASN132(OD1) |
| V126=LIG(C14)-LEU134(CD1) | V126=LIG(O20)-ASN132(ND2) |
| V127=LIG(C15)-ILE10(CG2) | V127=LIG(C21)-VAL18(CG1) |
| V128=LIG(C15)-ILE10(CD1) | V128=LIG(C21)-ASN132(ND2) |
| V129=LIG(C15)-HIE84(O) | V129=LIG(C21)-ASP145(CB) |
| V130=LIG(C15)-GLN85(O) | V130=LIG(C21)-ASP145(OD2) |
| V131=LIG(C15)-LEU134(CG) | V131=LIG(C22)-ILE10(CB) |
| V132=LIG(C15)-LEU134(CD1) | V132=LIG(C22)-ILE10(CG1) |
| V133=LIG(C15)-LEU134(CD2) | V133=LIG(C22)-ILE10(CD1) |
| V134=LIG(C16)-ILE10(CG2) | V134=LIG(C22)-LEU83(O) |
| V135=LIG(C16)-GLN85(O) | V135=LIG(C22)-GLN85(C) |
| V136=LIG(C16)-ASP86(CA) | V136=LIG(C22)-GLN85(O) |
| V137=LIG(C16)-ASP86(CB) | V137=LIG(C22)-ASP86(N) |
| V138=LIG(C16)-LEU134(CG) | V138=LIG(C22)-ASP86(CB) |
| V139=LIG(C16)-LEU134(CD2) | V139=LIG(C22)-ASP86(OD2) |
| V140=LIG(C17)-ILE10(CG2) | V140=LIG(C22)-LEU134(CD2) |
| V141=LIG(C17)-GLN131(O) | V141=LIG(C23)-VAL18(CB) |
| V142=LIG(C17)-LEU134(CG) | V142=LIG(C23)-VAL18(CG1) |
| V143=LIG(C17)-LEU134(CD1) | V143=LIG(C23)-VAL18(CG2) |
| V144=LIG(C17)-LEU134(CD2) | V144=LIG(C23)-ASP145(CB) |
| V145=LIG(C18)-ILE10(CG2) | V145=LIG(C24)-ILE10(CB) |
| V146=LIG(C18)-GLN131(O) | V146=LIG(C24)-ILE10(CG2) |
| V147=LIG(C18)-LEU134(CD2) | V147=LIG(C24)-GLN85(CA) |
| V148=LIG(C19)-ILE10(CG2) | V148=LIG(C24)-GLN85(C) |
| V149=LIG(C19)-ILE10(O) | V149=LIG(C24)-ASP86(N) |
| V150=LIG(C19)-ASP86(CB) | V150=LIG(C24)-ASP86(OD2) |
| V151=LIG(C19)-ASP86(CG) | V151=LIG(C25)-ILE10(CG1) |
| V152=LIG(C19)-ASP86(OD1) | V152=LIG(C25)-ILE10(CD1) |
| V153=LIG(C19)-ASP86(OD2) | V153=LIG(C25)-VAL18(CG2) |
| V154=LIG(C20)-LYS88(CB) | V154=LIG(C25)-ALA31(CB) |
| V155=LIG(C20)-LYS88(CG) | V155=LIG(C25)-LEU134(CG) |
| V156=LIG(C20)-GLN131(CA) | V156=LIG(C25)-LEU134(CD1) |
| V157=LIG(C20)-GLN131(CB) | V157=LIG(C25)-LEU134(CD2) |
| V158=LIG(C20)-GLN131(CG) | V158=LIG(C26)-ILE10(CG1) |
| V159=LIG(C20)-GLN131(C) | V159=LIG(C26)-ILE10(CD1) |
| V160=LIG(C20)-GLN131(O) | V160=LIG(C26)-VAL18(CG2) |
| V161=LIG(C21)-ILE10(O) | V161=LIG(C26)-LEU134(CG) |
| V162=LIG(C21)-GLY11(CA) | V162=LIG(C26)-LEU134(CD1) |
| V163=LIG(C21)-ASP86(OD1) | V163=LIG(C26)-LEU134(CD2) |
| V164=LIG(C22)-ILE10(CG2) | V164=LIG(C27)-VAL18(CB) |
| V165=LIG(C22)-ILE10(O) | V165=LIG(C27)-VAL18(CG1) |
| V166=LIG(C22)-ASP86(CA) | V166=LIG(C27)-VAL18(CG2) |
| V167=LIG(C22)-ASP86(CB) | V167=LIG(C27)-LEU134(CD2) |
| V168=LIG(C22)-ASP86(CG) | V168=LIG(C27)-ALA144(CB) |
| V169=LIG(C22)-ASP86(OD1) | V169=LIG(C28)-ILE10(CG1) |
| V170=LIG(C22)-ASP86(OD2) | V170=LIG(C28)-ILE10(CD1) |
| V171=LIG(C23)-ILE10(CG2) | V171=LIG(C28)-VAL18(CB) |
| V172=LIG(C23)-ILE10(O) | V172=LIG(C28)-VAL18(CG1) |
| V173=LIG(C23)-GLN85(C) | V173=LIG(C28)-VAL18(CG2) |
| V174=LIG(C23)-GLN85(O) | V174=LIG(C28)-LEU134(CD2) |
| V175=LIG(C23)-ASP86(N) | V175=LIG(C28)-ALA144(CB) |
| V176=LIG(C23)-ASP86(CA) | V176=LIG(C29)-LYS129(CE) |
| V177=LIG(C23)-ASP86(CB) | V177=LIG(C29)-LYS129(NZ) |
| V178=LIG(C23)-ASP86(CG) | V178=LIG(C29)-GLN131(CB) |
| V179=LIG(C23)-ASP86(OD1) | V179=LEU(CA)-LYS20(C) |
| V180=LIG(C23)-ASP86(OD2) | V180=GLN(C)-VAL18(C) |
| V181=LIG(C23)-LEU134(CD2) | V181=LEU(C)-1011.086(ILE) |
| V182=LEU(CA)-LYS20(C) |  |
| V183=GLN(C)-VAL18(C) |  |
| V184=LEU(C)-1011.086(ILE) |  |

### 6. ML Training Results and Feature Analysis

#### Analytical model system

We generated 180 independent trajectories of 500 steps each and for each trajectory frame we calculated the 180 input features. We split the dataset into a training set (70% of all trajectories) and a test set (30% of all trajectories) for the ML training. In addition, we run additional 50 independent simulations for validation of the extent of any overfitting. We determined the ML prediction accuracy at different times by varying total number of frames (i.e., simulation intervals) and starting from different initial frames throughout the trajectories.

**Figure S5.**
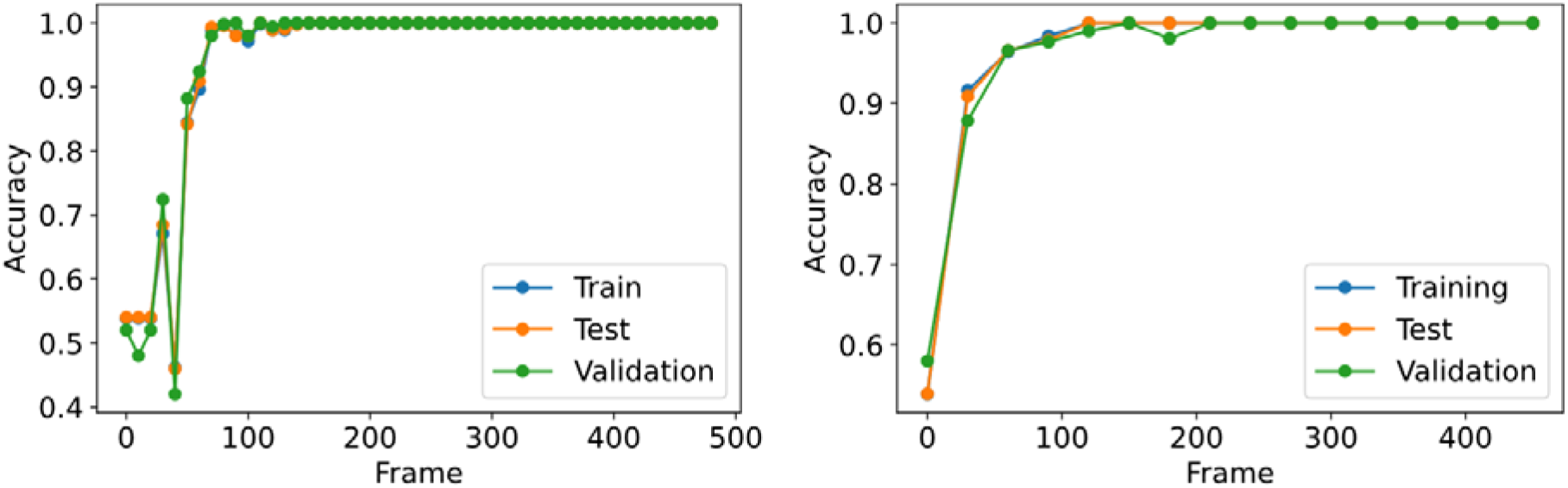
Accuracy of the Multi-Layer Perceptron (MLP) prediction for the dataset with 1 DW and 24 SW potentials at different starting times using 10 frame intervals (left) and 30 frame intervals (right).

For the result that a time range of 30 steps starting from the 30^th^ step (from the 30^th^ to 60^th^ step) to be satisfactory for this approach. Thus, we started to train on that time range 100 models for both GBDT and MLP to proceed with our feature analysis. Some average values for accuracy and training epochs are shown in the following table:

**Table S3.** Table containing the average accuracies (for training, test and validation) and number of epochs used for training of GBDT and MLP methods over the 100 independent replicates of our procedure, for both types of datasets (1 DW and the 5 DW potentials) tested.

|  | ⟨Training Accuracy (%)⟩ | ⟨Test Accuracy (%)⟩ | ⟨Validation Accuracy (%)⟩ | ⟨Epochs⟩ |
| --- | --- | --- | --- | --- |
| GBDT (1DW) | 100.00 | 99.72 | 91.45 | 500 |
| MLP (1DW) | 94.83 | 94.73 | 93.04 | 204 |
| GBDT (5DW) | 100.00 | 99.80 | 91.64 | 500 |
| MLP (5DW) | 91.85 | 91.83 | 89.32 | 311 |

#### Analytical model with 5 - Double Well potentials MLTSA vs GBDT

**Figure S6.**
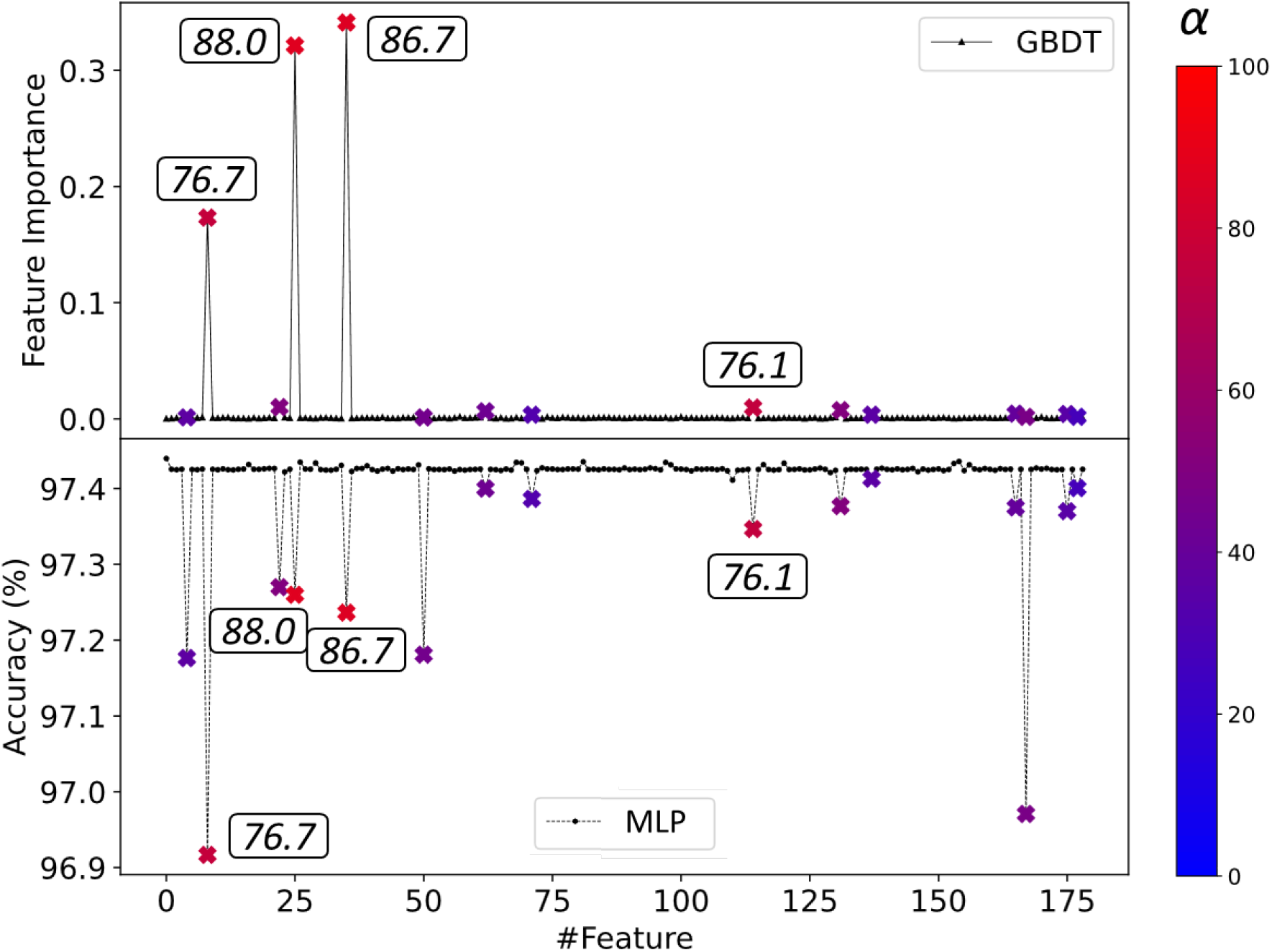
Comparison between GBDT (top) and MLTSA (bottom) feature analysis methods and the captured correlated features on the 5DW dataset for the analytical model. From blue (0%) to red (100%), the correlated features are marked depending on the level of correlation. The top four features are labelled. The baseline symbols (black) in the top plot show the feature importance values for each feature which are close to 0 when irrelevant. The baseline symbols (black) at the bottom plot show the accuracy of the trained data when each feature is altered with its global mean across simulations.

#### CDK2 System

Details of the trained models during the MLTSA using 0.1, 0.15, 0.3, 0.5, 0.75, 1, 1.5, 3, and the full 5 ns length of the downhill trajectories for each system (4fkw and 3sw4) are listed below. In addition to testing the different lengths of trajectories, the percentage of data to use from the latter end of the trajectory at each time frame (i.e., the 50% latter end of 0.1 ns would correspond to data from 0.05 ns to 0.1 ns) was also tested. The number of simulations available and the number of epochs until convergence for each model are also listed, as well as their accuracy on a set of independent simulations (Validation set). This set is comprised of the 25% of the available data from 4fkw and 3sw4 having 35 and 37 simulations to test the accuracy, respectively.

**Table S4.I.** The table below comprises the details of the models tested on 4fkw data as well as their accuracy on the validation set. The first column corresponds to the time frame of trajectory data used from the beginning. The data column corresponds to the percentage of latter simulation time used to train each model. The third column has the number of epochs until convergence of the model and the last column shows the accuracy on the validation set.

| System |  | Simulations |  |
| --- | --- | --- | --- |
| 4fkW |  | 139 |  |
| Time (ns) | Data | Epochs | Accuracy |
| 0.1 | 5% | 198 | 48% |
|  | 10% | 142 | 48% |
|  | 25% | 180 | 47% |
|  | 50% | 190 | 47% |
| 0.15 | 5% | 166 | 45% |
|  | 10% | 211 | 49% |
|  | 25% | 223 | 47% |
|  | 50% | 221 | 45% |
| 0.3 | 5% | 174 | 63% |
|  | 10% | 165 | 63% |
|  | 25% | 236 | 59% |
|  | 50% | 205 | 54% |
| 0.5 | 5% | 173 | 53% |
|  | 10% | 210 | 55% |
|  | 25% | 295 | 46% |
|  | 50% | 358 | 55% |
| 0.75 | 5% | 179 | 45% |
|  | 10% | 112 | 55% |
|  | 25% | 304 | 53% |
|  | 50% | 206 | 52% |
| 1 | 5% | 105 | 52% |
|  | 10% | 170 | 50% |
|  | 25% | 248 | 55% |
|  | 50% | 333 | 54% |
| 1.5 | 5% | 131 | 61% |
|  | 10% | 195 | 63% |
|  | 25% | 338 | 63% |
|  | 50% | 354 | 62% |
| 3 | 5% | 148 | 81% |
|  | 10% | 110 | 81% |
|  | 25% | 151 | 79% |
|  | 50% | 201 | 74% |
| 5 | 5% | 19 | 100% |
|  | 10% | 18 | 99% |
|  | 25% | 22 | 93% |
|  | 50% | 73 | 87% |

**Table S4.II.** The table below comprise the details of the models tested on 3sw4 data as well as their accuracy on the validation set. The first column corresponds to the time frame of trajectory data used from the beginning. The data column corresponds to the percentage of latter simulation time used to train each model. The third column has the number of epochs until convergence of the model and the last column shows the accuracy on the validation set.

| System |  | Simulations |  |
| --- | --- | --- | --- |
| 3sw4 |  | 148 |  |
| Time (ns) | Data | Epochs | Accuracy |
| 0.1 | 5% | 192 | 57% |
|  | 10% | 310 | 63% |
|  | 25% | 239 | 53% |
|  | 50% | 194 | 53% |
| 0.15 | 5% | 279 | 67% |
|  | 10% | 215 | 68% |
|  | 25% | 175 | 71% |
|  | 50% | 293 | 70% |
| 0.3 | 5% | 130 | 57% |
|  | 10% | 123 | 66% |
|  | 25% | 293 | 53% |
|  | 50% | 353 | 55% |
| 0.5 | 5% | 236 | 62% |
|  | 10% | 210 | 62% |
|  | 25% | 333 | 63% |
|  | 50% | 191 | 63% |
| 0.75 | 5% | 231 | 66% |
|  | 10% | 249 | 69% |
|  | 25% | 320 | 65% |
|  | 50% | 217 | 62% |
| 1 | 5% | 292 | 62% |
|  | 10% | 220 | 60% |
|  | 25% | 249 | 63% |
|  | 50% | 229 | 66% |
| 1.5 | 5% | 229 | 67% |
|  | 10% | 285 | 71% |
|  | 25% | 277 | 62% |
|  | 50% | 189 | 62% |
| 3 | 5% | 246 | 82% |
|  | 10% | 342 | 75% |
|  | 25% | 204 | 73% |
|  | 50% | 153 | 73% |
| 5 | 5% | 34 | 100% |
|  | 10% | 38 | 99% |
|  | 25% | 103 | 97% |
|  | 50% | 83 | 87% |

**Figure S7.**
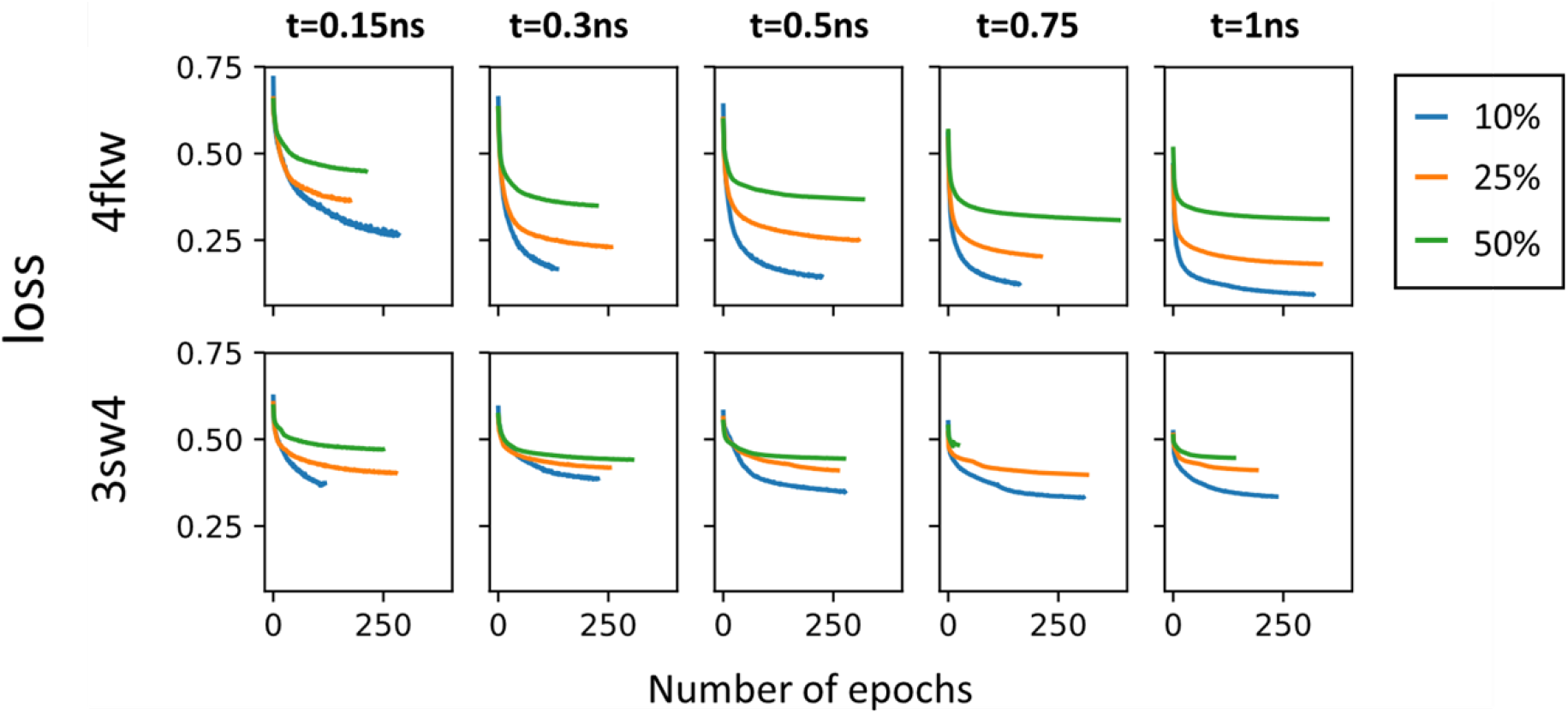
Plots of the loss function evolution through the training epochs at different time frames with different percentages of data from the end for both 3sw4 (18K ligand) and 4fkw (62K ligand) CDK2 systems.

To understand the relationship between the accuracy of predictions and the data used to make those predictions, we trained the MLP with several different datasets. As described in the main text, each trajectory provided a set of distances from the simulated trajectory at particular timeframes, and each dataset was made up of a set of such timeframe elements. The trainings used different timeframes of the trajectories: at 0.3, 0.5, 0.75, 1, 1.5, 3 and 5 ns. For each of these datasets we calculated the accuracy of the predictions for each of the three systems. The models provide good accuracy from the very initial frames of the simulations. For example, at 0.1 ns we have an accuracy of 79.5% for ligand 18K and 83.6% for 62K.

**Figure S8.**
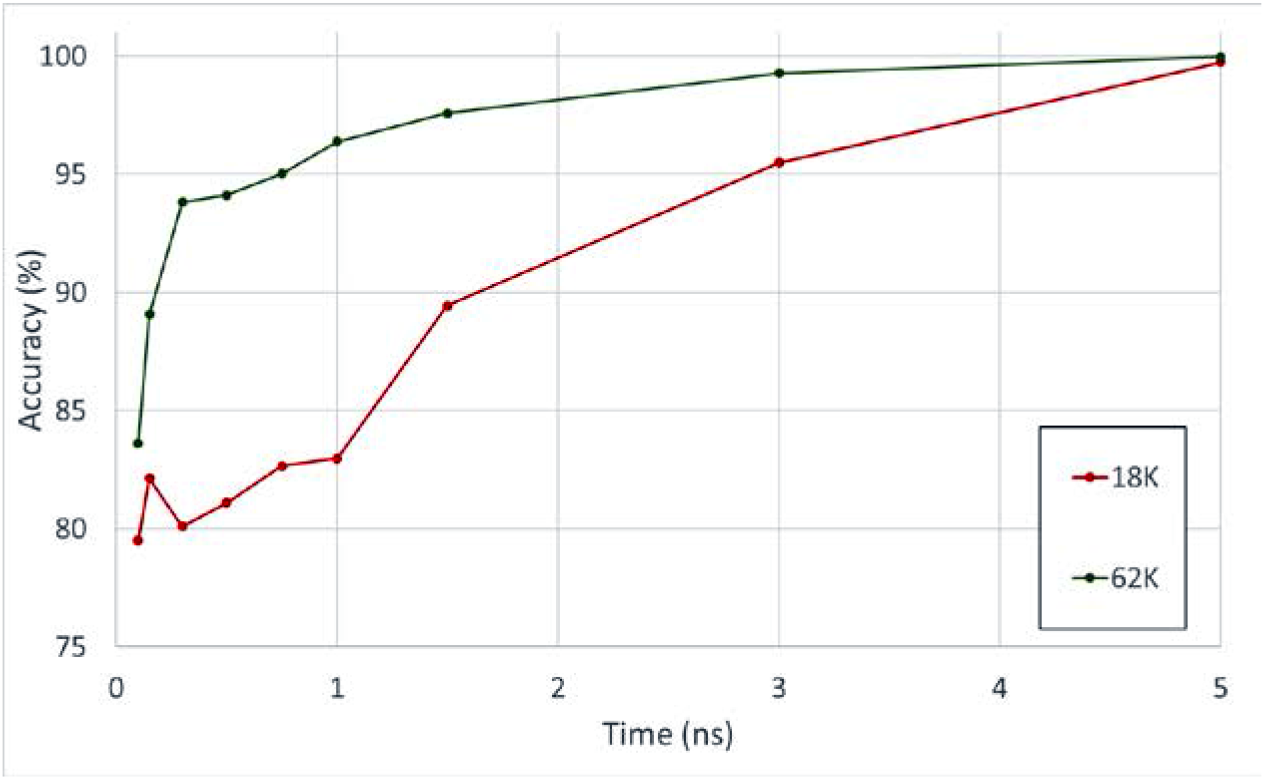
ML accuracy prediction at different time frames using MLP for 18K in red and 62K in green.

### 7. Free Energy Profiles

For each system, we performed three independent replicas. The PMF is plotted along the string windows. For each replica, the number of distances included in the string depends on the unbinding trajectory. The number of distances used in each system are given in Table S2. (Fig. S2.I-S2.VI above for details).

**Figure S9.**
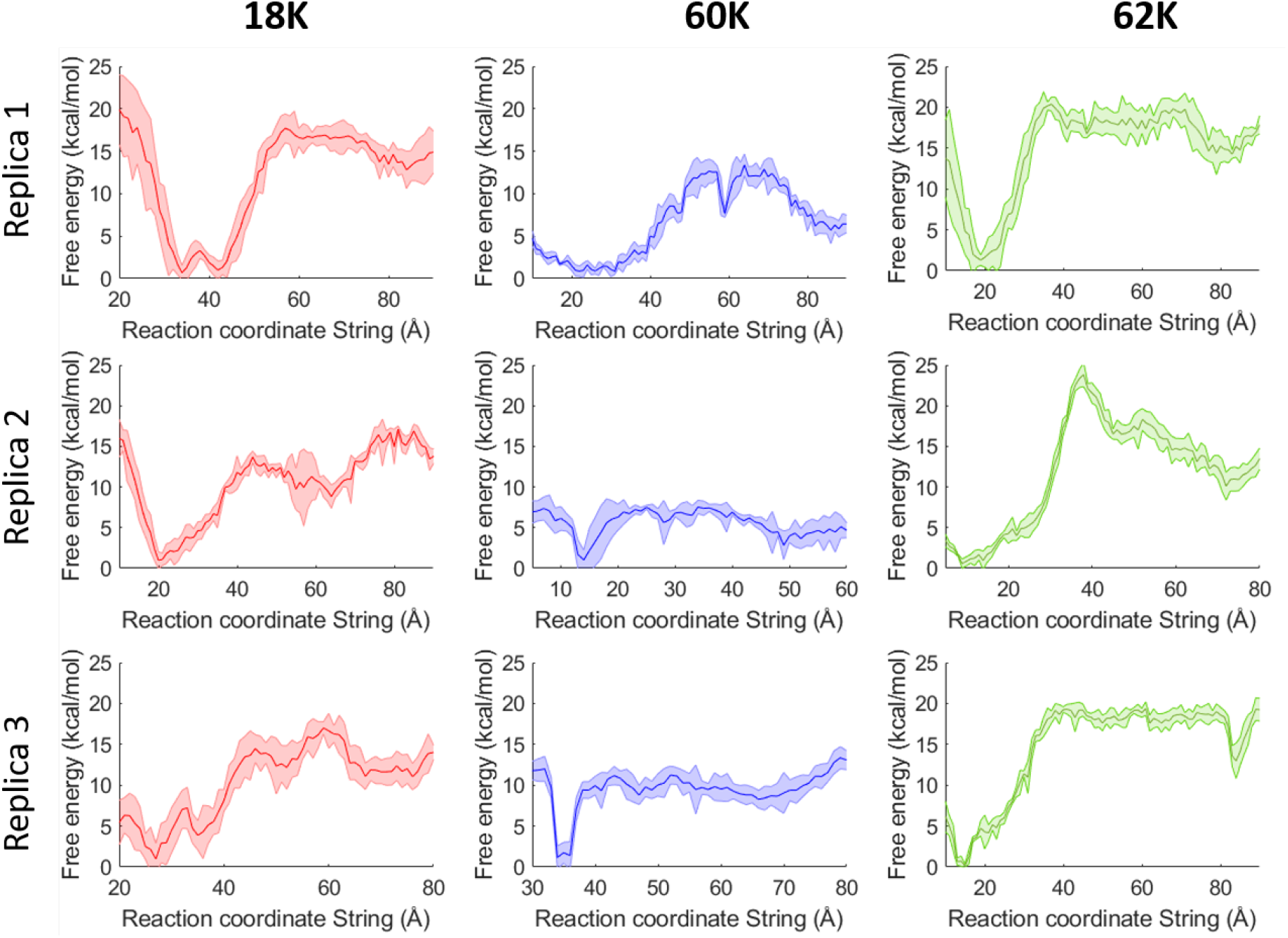
PMF of the unbinding path for 18K, 60K and 62K. The standard error shown as shaded area are obtained by dividing the full dataset into 4 subgroups.

**Table S5.** Number of distances included in the RC for each replica.

| System | Number of distances |
| --- | --- |
| 18K Replica 1 | 21 |
| 18K Replica 2 | 38 |
| 18K Replica 3 | 33 |
| 62K Replica 1 | 46 |
| 62K Replica 2 | 38 |
| 62K Replica 3 | 43 |

### 8. 60K/4FKU system

An additional ligand was tested with our unbinding approach, an oxindole carboxylic acid derivative (60K) based on the 4fku structure (Fig. S10). The unbinding procedure was carried out as described for the other ligands. After performing the string calculations for the 4fku system, 60K presented a change in conformation, more specifically, a cis-trans conversion of the hydrazineyl N=C bond (Fig. S11). This conformational change would only be expected at very high energy costs, and it is a combined artifact of the force field and the biasing procedure. The Z (cis) to E (trans) conversion allowed the 60K ligand to unbind with a significantly lower free energy barrier than its analogue, 4fkw (Figs. S12 and S13). They both share a dihedral angle (φ), which corresponds to this transformation, defined between atoms N6-N9-C14-C16 for 60K and N1-N3-C25-C26 for 62K (Fig. S12). On one hand, this is partly due to the initially strong constraints from the string method that can be corrected in the future. On the other hand, this is also due to the too low energy of the trans form and the too low barrier for the isomerisation as compared to the DFT calculations (Fig. S13). Tas a result, the final unbinding free energy barrier (Fig. S9, middle, blue) is ∼10 kcal/mol lower than the experimental (20.01 (±0.12) kcal mol^-1^) value for all three replicas (9.96 (±1.5) kcal mol^-1^).

**Figure S10.**
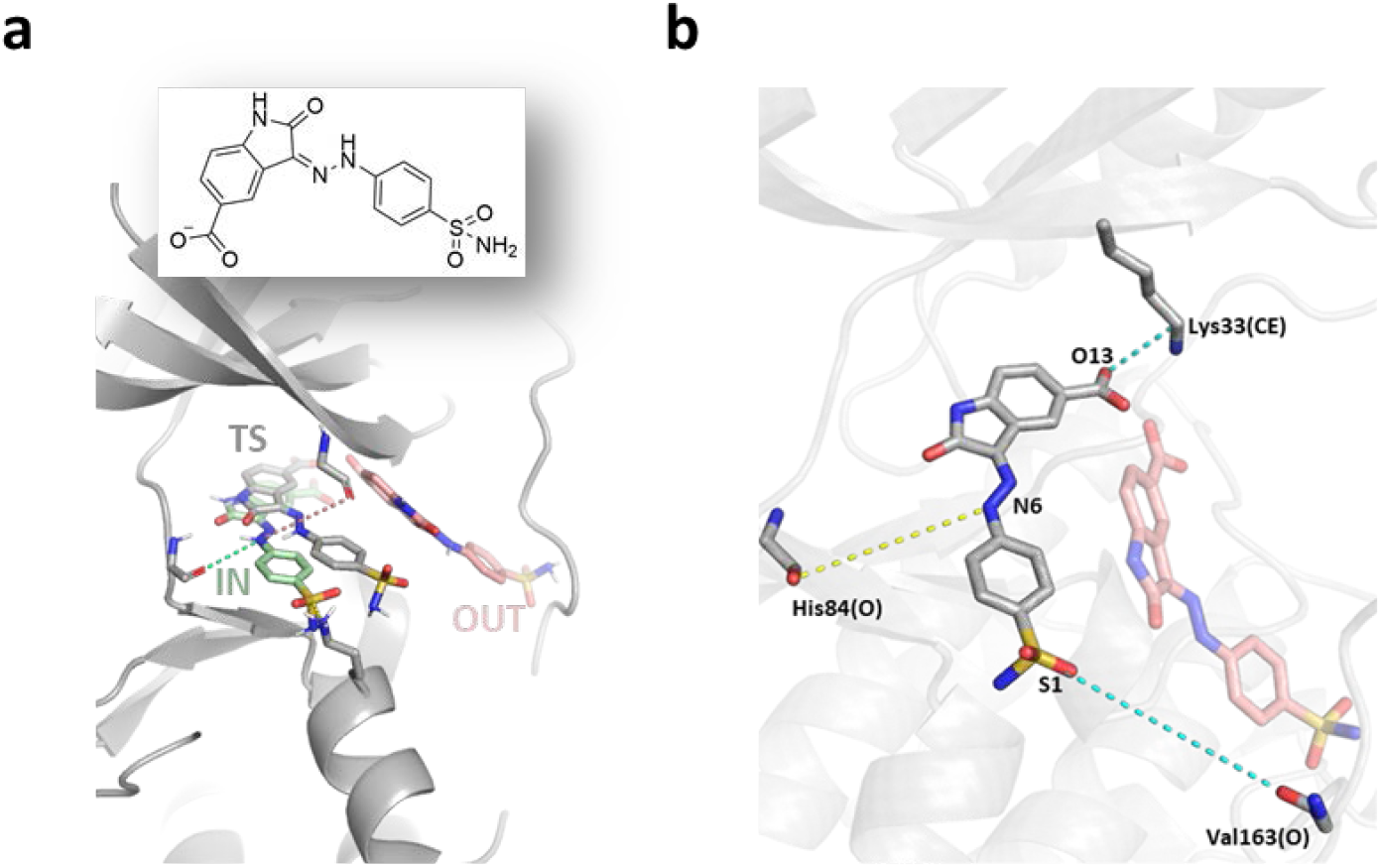
Left (a): CDK2 bound to 60K, the chemical structure of the ligand oxindole carboxylic acid derivative is drawn in the inset. Bound state (IN) originated from PDB structure 4fku. Structural details of the ATP pocket are shown with the ligand in the bound state (green), unbound (red) and transition state (grey). Right (b): common CVs obtained from the unbinding replicas of 60K, representative distances are shown in dashed lines (yellow: interaction from the initial structure, cyan: interaction found during the unbinding trajectory), red sticks represent the ligand when it is outside the pocket. The displayed distances appear in all three replicas for 60K.

**Figure S11.**
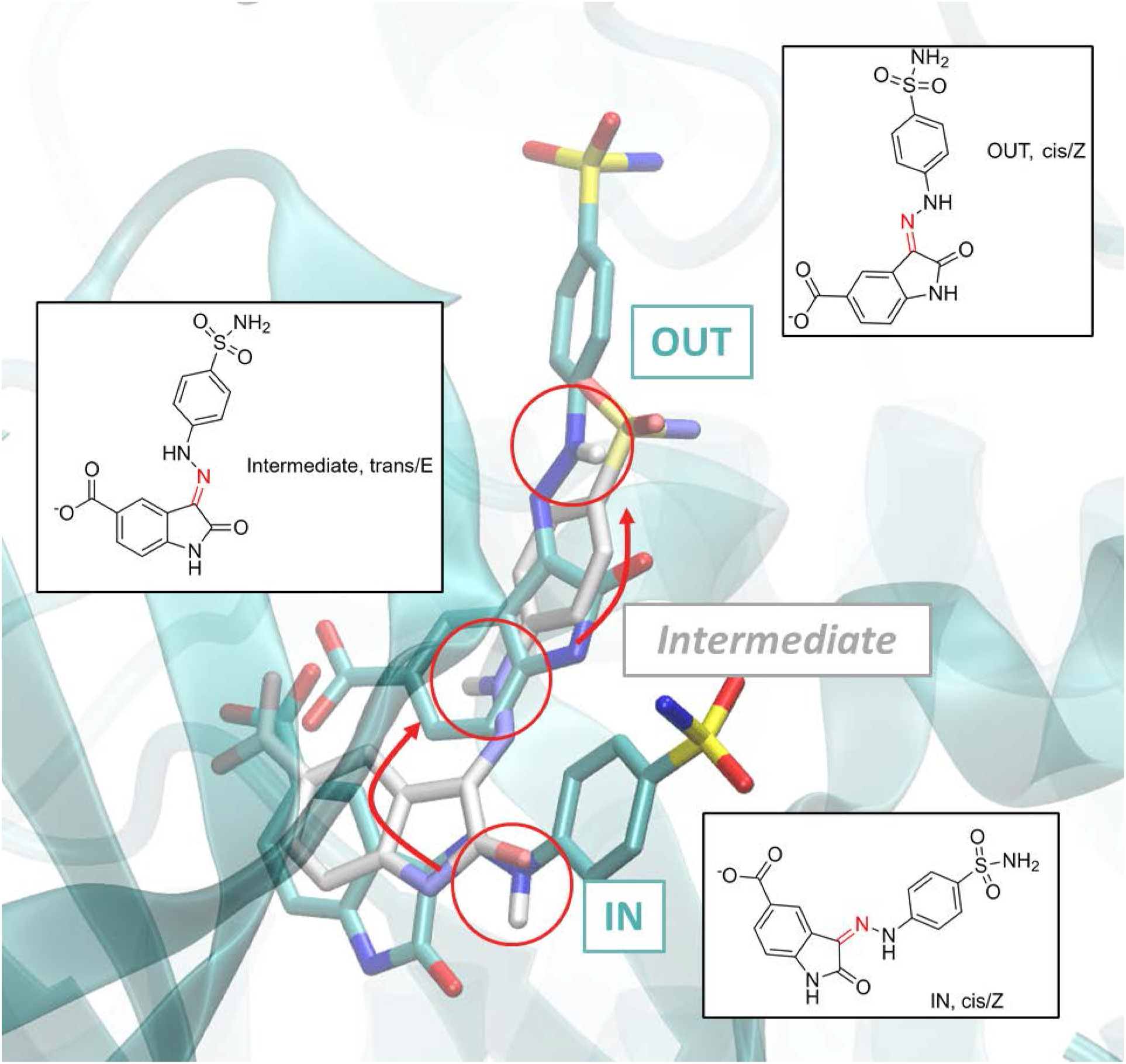
60K ligand structures within the CDK2 binding pocket from three different umbrella windows portraying the cis-trans conversion through the unbinding pathway from IN (cis/Z, red circle) via the intermediate (trans/E, red circle, white sticks) to OUT (cis/Z, red circle) structures.

**Figure S12.**
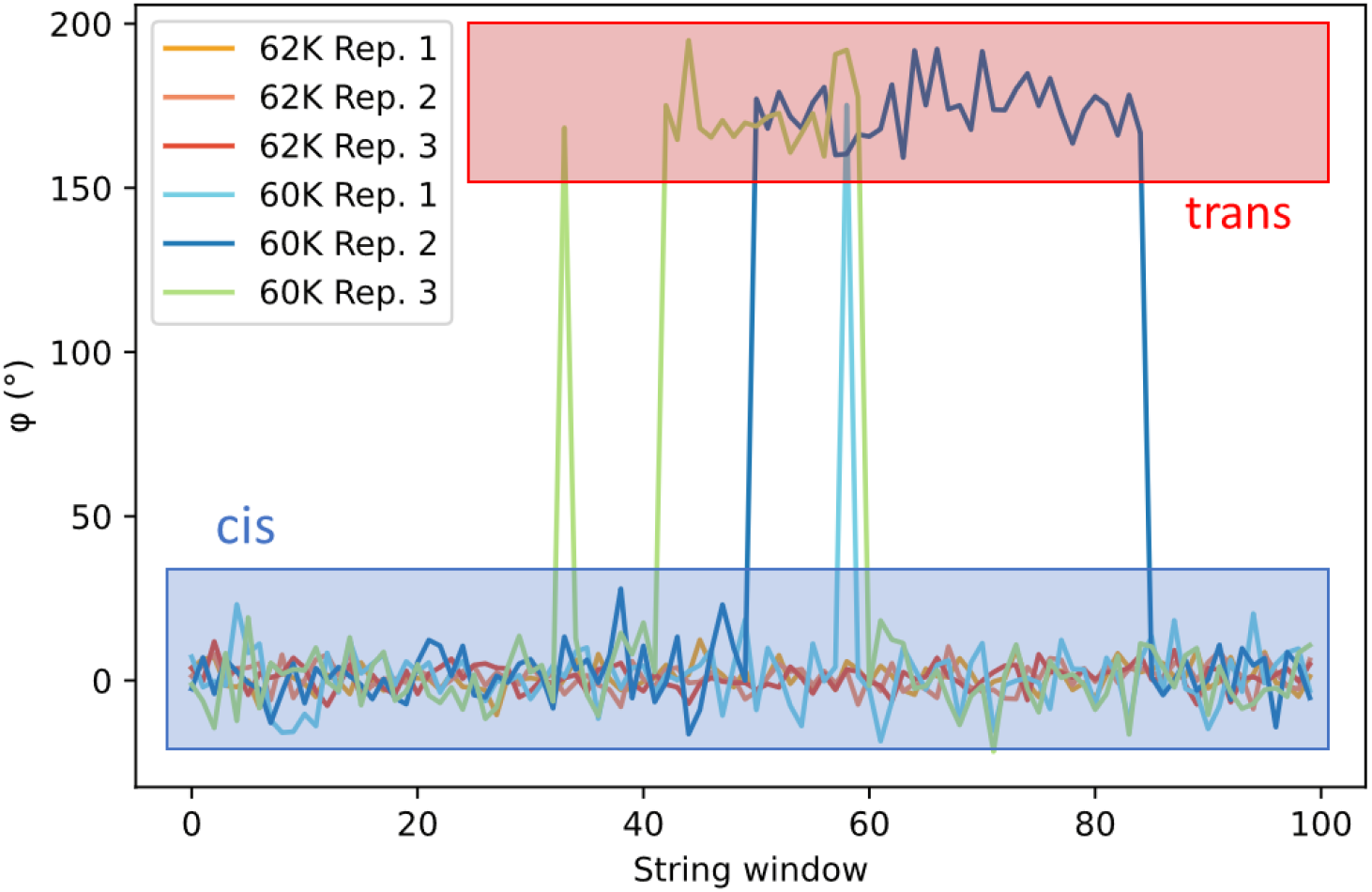
Values of the dihedral angles (φ) for both 4FKU (60K) and 4FKW (62K) throughout the umbrella sampling windows of all three replicas. φ is defined as the dihedral angle between atoms N6-N9-C14-C16 for 60K and C25-C26-N1-N3 for 62K. The conformations at φ∼0° correspond to the cis isomers and at φ∼180° correspond to the trans isomers.

**Figure S13.**
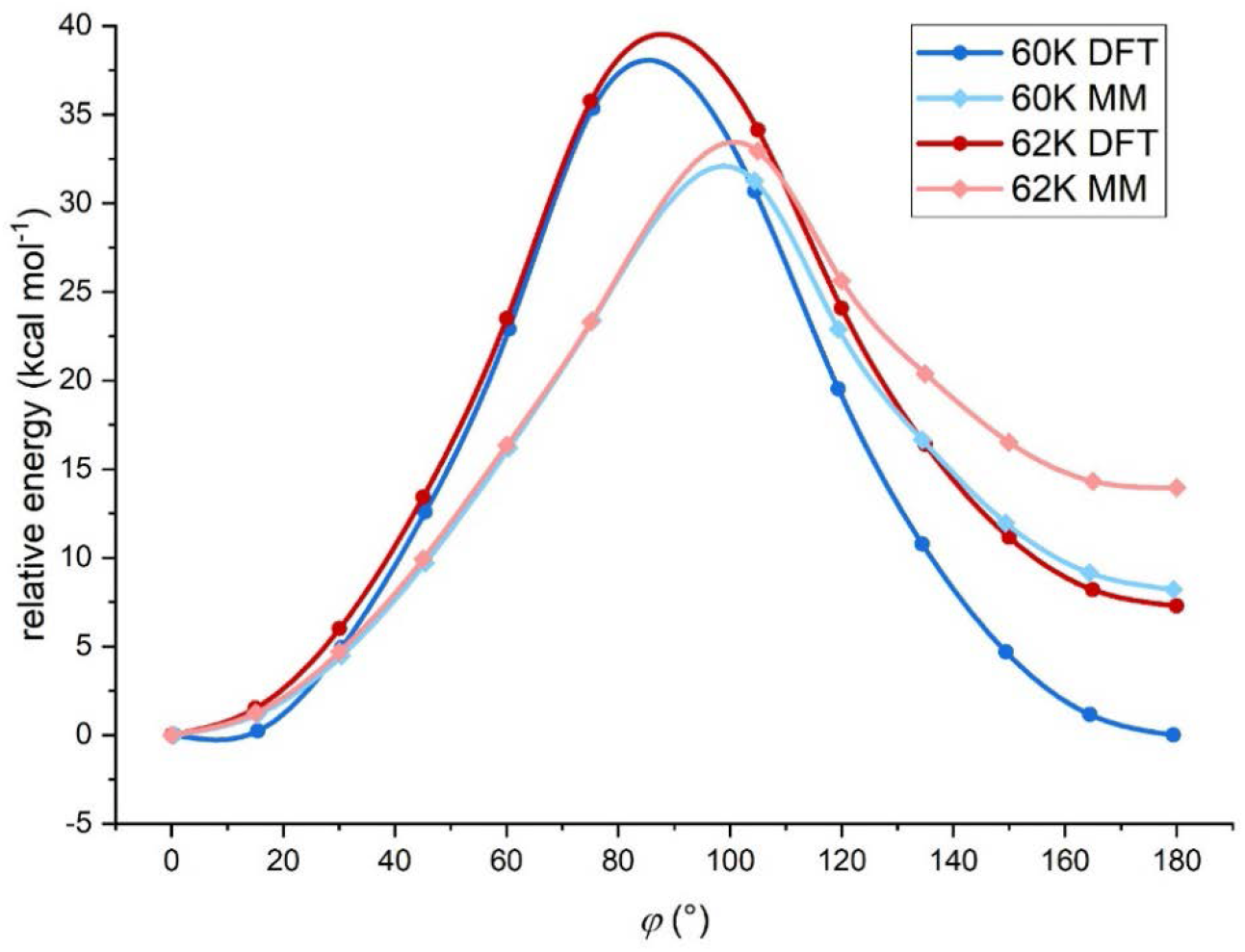
Relative energy values for 60K and 62K calculated at DFT (dark blue and dark red circles, respectively) and MM (light blue and pink squares, respectively) levels of theory for the cis-trans interconversion based on DFT optimized geometries along φ. The dihedral angle (φ) is defined between atoms N6-N9-C14-C16 for 60K and N1-N3-C25-C26 for 62K. The rotational barriers are lower in the force field (MM) than calculated at the DFT level. Note that the MM relative energy of the trans isomer with respect to the cis for 62K is about 10 kcal/mol higher than for 60K, contributing to the different behavior observed between the two similar ligands.

### 9. Validation of the ML Analysis

Figures S14.I and S14.II compare the results of our training against a simple binary classification model which attempts to classify the outcome as IN/OUT based on the CV values at a specific time (0.15, 0.3 and 0.5 ns). The dots show the CV values (as a sum of two key distances from Table S2.I) and are colored according to their outcome, red as OUT and green as IN. We then calculate the accuracy of the binary prediction at different thresholds represented by the black bars to obtain the highest possible accuracy using a single cutoff value (blue arrow). We compare these with the values obtained from the MLP (blue data, top of Figs. S14.I-II) and the GBDT (yellow data, top of Figs. S14.I-II).

**Figure S14.I.**
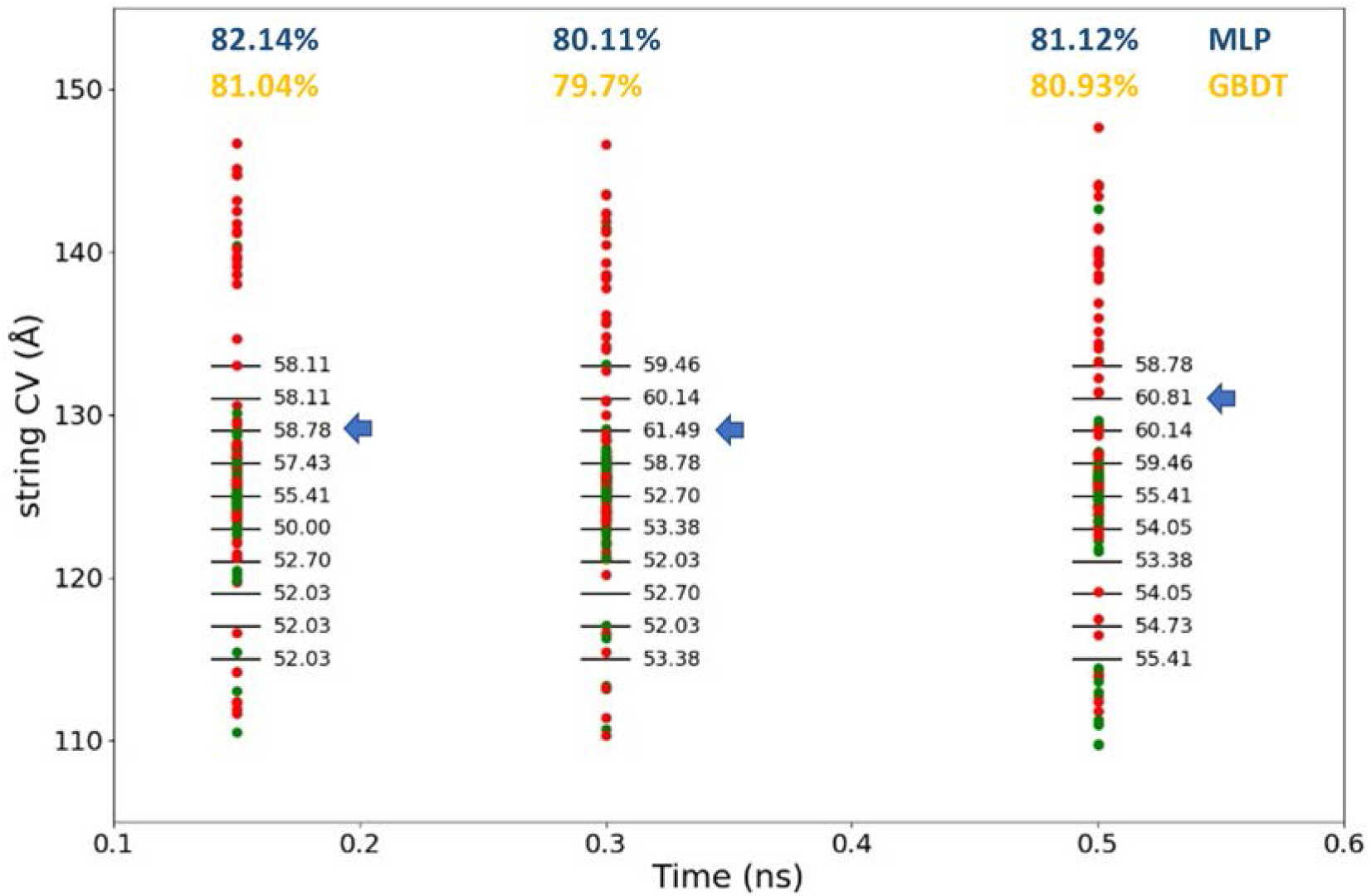
Comparison of the accuracy obtained from our MLTSA training (blue data) and GBDT (yellow data) with a simple binary classification model for ligand 18K at 0.15, 0.3 and 0.5 ns. Data points corresponding to different trajectories show the actual value of the string CV for IN (green) and OUT (red) trajectories.

**Figure S14.VII.**
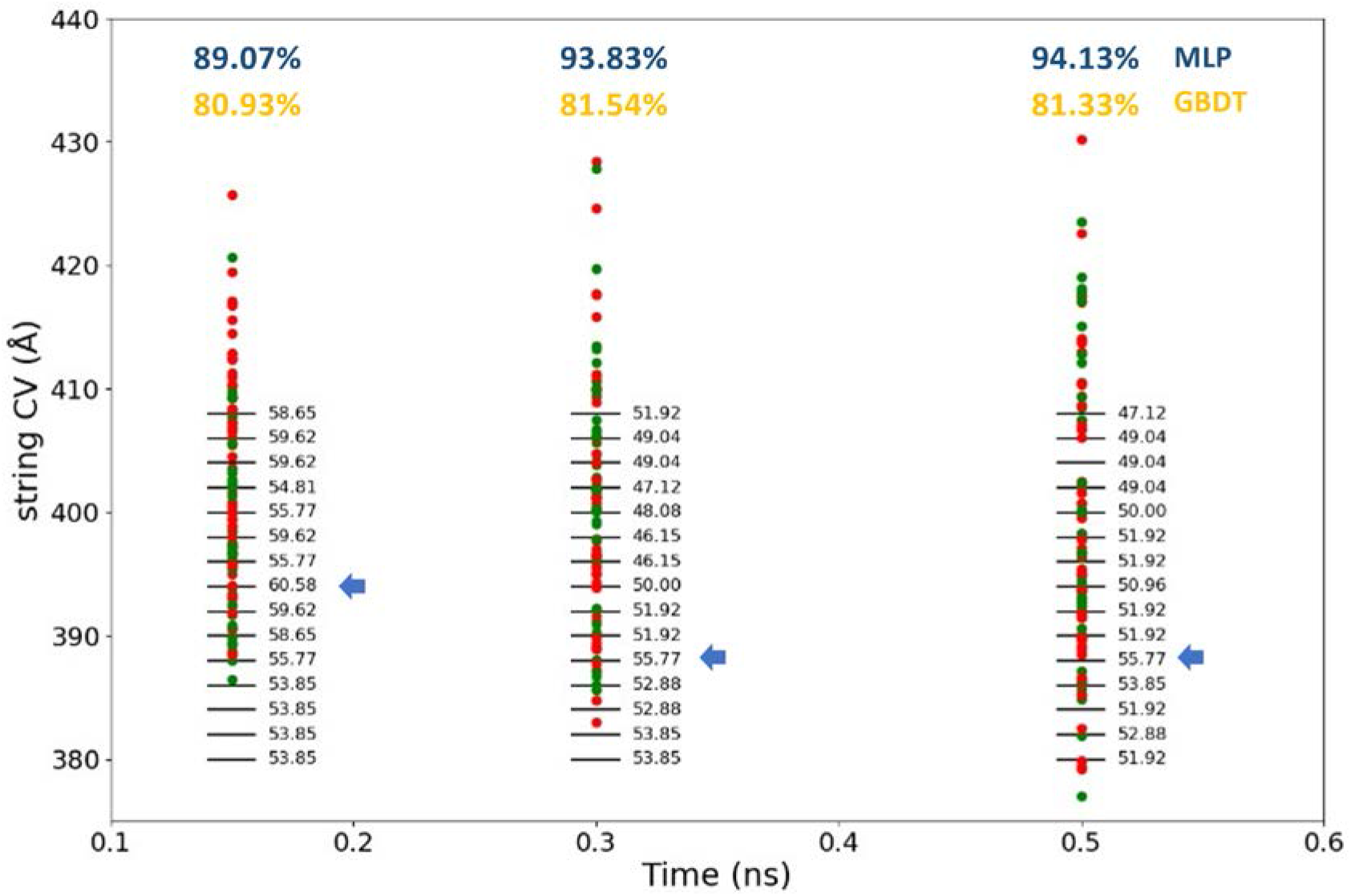
Comparison of the accuracy obtained from our MLTSA training (blue data) and GBDT (yellow data) with a simple binary classification model for ligand 62K at 0.15, 0.3 and 0.5 ns. Data points corresponding to different trajectories show the actual value of the string CV for IN (green) and OUT (red) trajectories.

### 10. Gradient Boosting Decision Trees

We used GBDT as an alternative approach to the MLP. The model was trained using the same amount of data fed for the MLP. We compared the results obtained from the MLTSA against the feature importances given by the GBDT. Overall, features resulting important from the MLTSA are also present in the GBDT, however, depending on the system we analyzed, additional important features were also detected from the GBDT’s important features. This suggests that the more complex non-linear behavior might lead to different performances for GBDT and the MLP as compared to the analytical model system.

**Figure S15.VIII.**
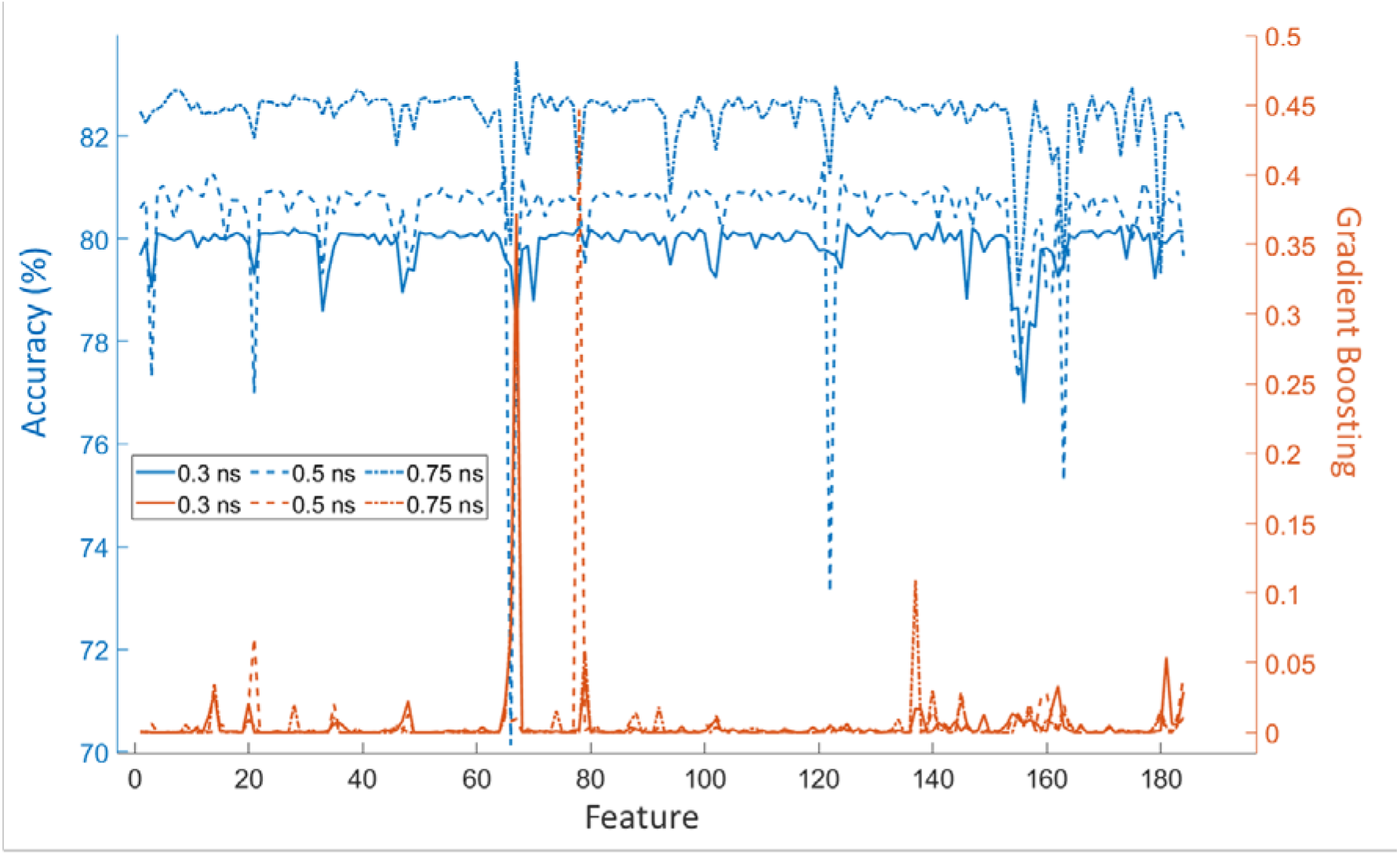
Comparison between GBDT feature importance (orange) and MLTSA accuracy drops (blue) at different times for the three systems for ligand 18K.

**Figure S15.IX.**
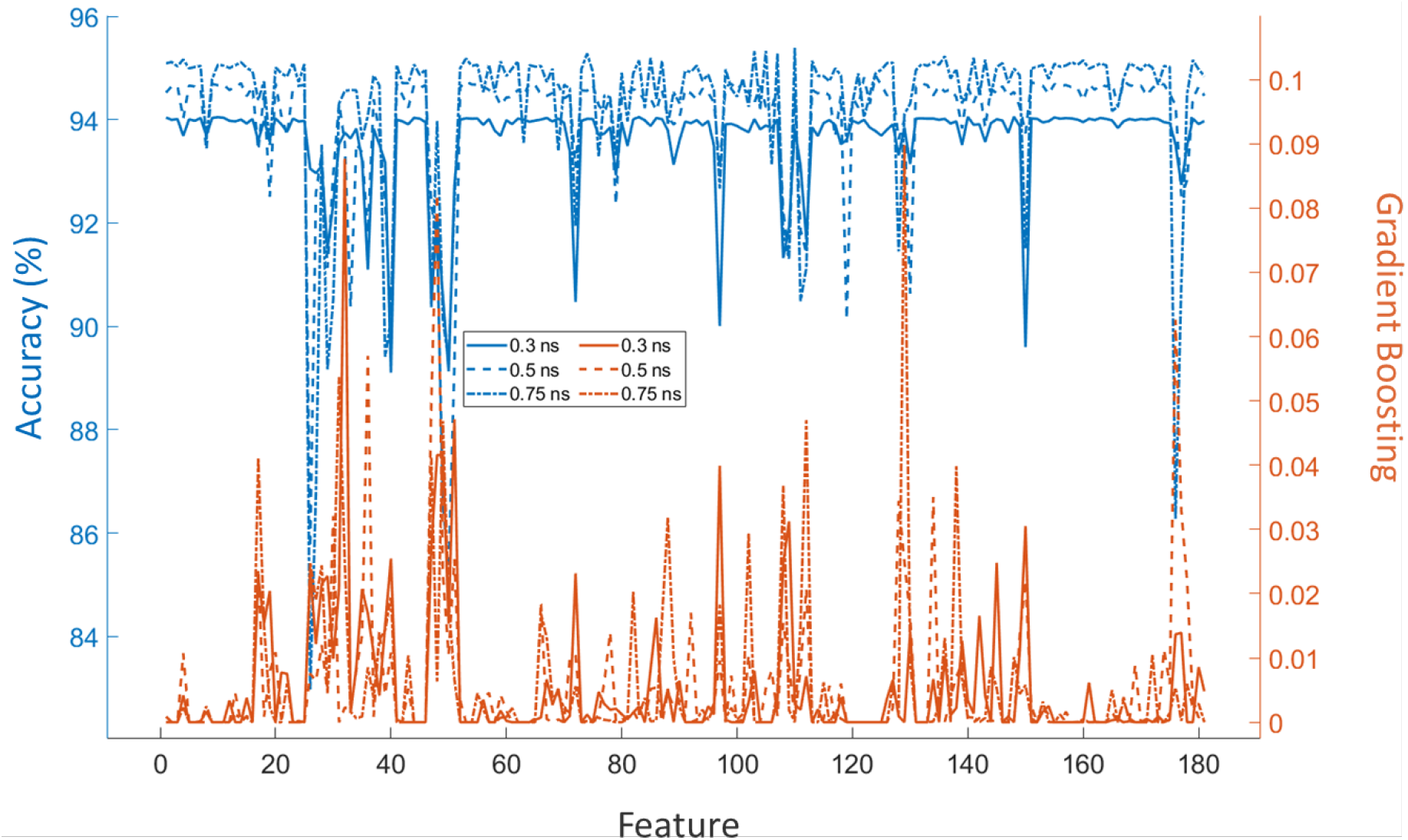
Comparison between GBDT feature importance (orange) and MLTSA accuracy drops (blue) at different times for the three systems for ligand 62K.

### 11. Additional resources

#### Animated trajectories

Animated GIF files showing the string trajectories for the three systems (3sw4, 4fku and 4fkw) of all replicas are available at the GitHub repository:

- https://github.com/pedrojuanbj/MLTSA-V1

#### Software package

A Python package of the analytical MLTSA example and corresponding Python code is accessible under the Python Package Index (PyPi) database:

- https://pypi.org/project/MLTSA/

#### Jupyter Notebook examples

Fully annotated *Jupyter Notebook* examples on how to apply the MLTSA approach for the analytical model among others, are available under the “*MLTSA_examples*” folder on our <u>GitHub</u> repository as well as when installing our Python <u>package</u>.

